# The integration of visual and target signals in V4 and IT during visual object search

**DOI:** 10.1101/370049

**Authors:** Noam Roth, Nicole C. Rust

**Affiliations:** Department of Psychology, University of Pennsylvania, Philadelphia, PA USA

**Keywords:** object search, visual attention, top-down signals, invariant object recognition, ventral visual pathway

## Abstract

Searching for a specific visual object requires our brain to compare the items in view with a remembered representation of the sought target to determine whether a target match is present. This comparison is thought to be implemented, in part, via the combination of top-down modulations reflecting target identity with feed-forward visual representations. However, it remains unclear whether top-down signals are integrated at a single locus within the ventral visual pathway (e.g. V4) or at multiple stages (e.g. both V4 and inferotemporal cortex, IT). To investigate, we recorded neural responses in V4 and IT as rhesus monkeys performed a task that required them to identify when a target object appeared across variation in position, size and background context. We found non-visual, task-specific signals in both V4 and IT. To evaluate whether V4 was the only locus for the integration of top-down signals, we evaluated several feed-forward accounts of processing from V4 to IT, including a model in which IT preferentially sampled from the best V4 units and a model that allowed for nonlinear IT computation. IT task-specific modulation was not accounted for by any of these feed-forward descriptions, suggesting that during object search, top-down signals are integrated directly within IT.

**NEW & NOTEWORTHY:** To find specific objects, the brain must integrate top-down, target-specific signals with visual information about objects in view. However, the exact route of this integration in the ventral visual pathway is unclear. In the first study to systematically compare V4 and IT during an invariant object search task, we demonstrate that top-down signals found in IT cannot be described as being inherited from V4, but rather must be integrated directly within IT itself.

## INTRODUCTION

Finding a sought object requires our brains to perform at least two non-trivial computations. First, we must determine the identities of the objects in view, across variation in details such as their position, size, and background context. Second, we must compare this visual representation (of what we are looking at) with a remembered representation (of what we are looking for) to determine whether our target is in view. Considerable evidence suggests that computations in the primate ventral visual pathway, including brain areas V1, V2, V4 and IT, support the process of invariant object recognition (reviewed by DiCarlo et al. 2012). Within V4 and IT, many neurons are also modulated by information about target identity as well as whether an image is a target match (Haenny et al. 1988; Maunsell et al. 1991; Eskandar et al. 1992; Leuschow et al. 1994; Gibson and Maunsell 1997; Chelazzi et al. 1998; Chelazzi et al.2001; Bichot et al. 2005; Pagan et al. 2013; Kosai et al. 2014; Roth and Rust 2018a). However, the route by which these signals arrive in V4 and IT remains unclear.

Here we present two proposals for how top-down signals reflecting the identity of a sought target and/or whether the object in view is a target match might arrive within V4 and IT during object search. In the first proposal (Fig 1a), V4 serves as the sole locus of the combination of visual and top-down information, and IT receives this information via feed-forward propagation from V4. In the second (Fig 1b), top-down information is integrated directly in IT, either exclusively or in addition to its integration in V4. A number of studies report that task-relevant signals increase in a gradient-like fashion across the early visual hierarchy (i.e. V1, V2 and V4) during covert spatial attention and feature-based attention tasks (reviewed by Noudoost et al. 2010), consistent with the integration of top-down signals at multiple, early stages. However, the few studies that have compared top-down modulation at higher stages of the pathway, including V4 and IT, report it to be matched, both during visual target search (Chelazzi et al. 1998; Chelazzi et al. 2001) as well as one covert spatial attention task (Moran and Desimone 1985). Additionally, while one might postulate that because receptive fields are smaller in V4, the brain would be more likely to integrate top-down signals in IT when a task requires spatial invariance (such as detecting the change in a feature despite its position), V4 feature-based attention effects have in fact been demonstrated to extend globally across the visual field (reviewed by Maunsell and Treue 2006; Cohen and Maunsell 2011). Together, the published evidence is thus more supportive of the proposal presented in Fig 1a, where IT inherits its visual as well as top-down information exclusively from V4.

**Figure 1.**
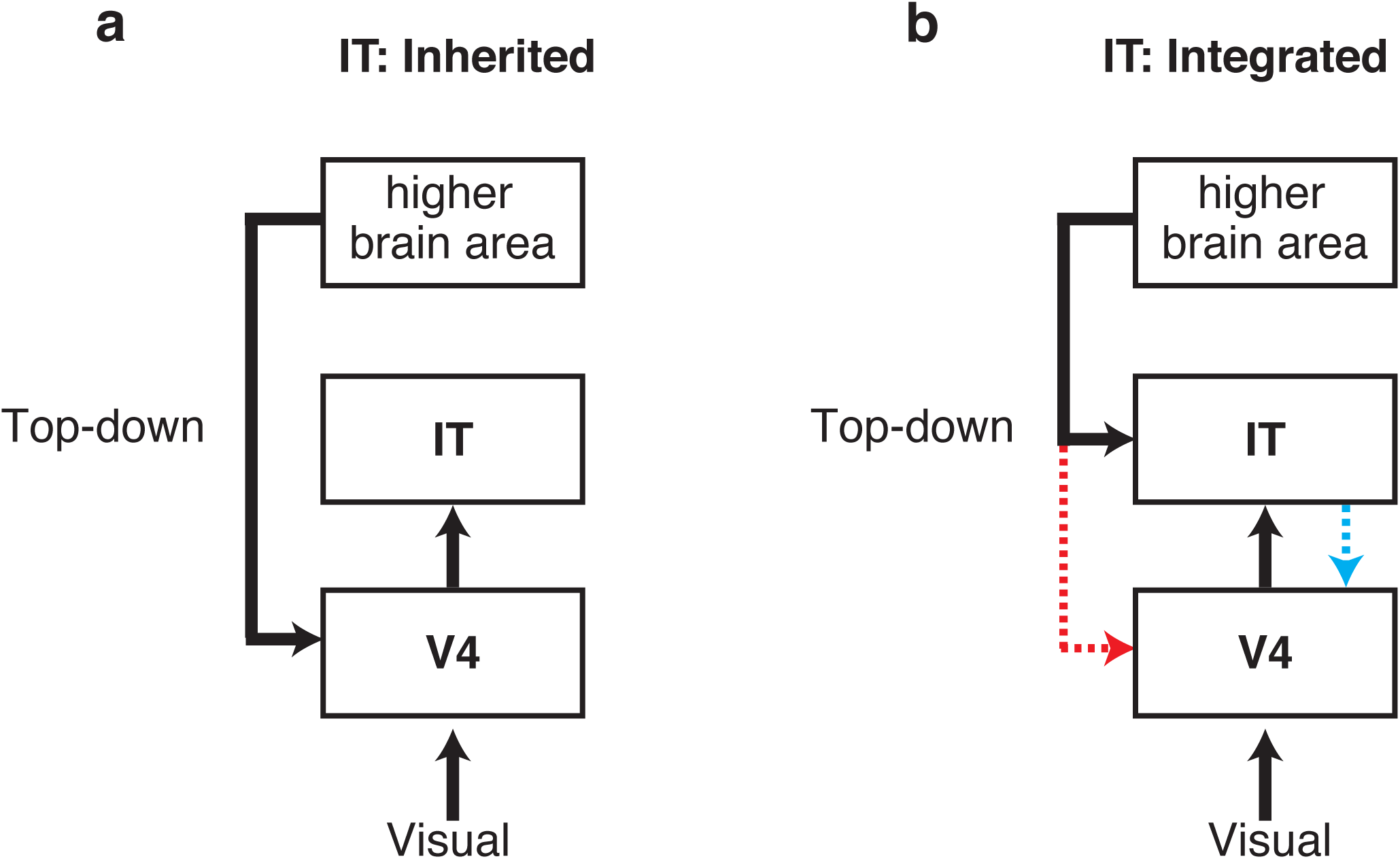
Proposals for how top-down information might arrive within V4 and IT during object search. **a)** The class of “IT: inherited” proposals predict that top-down information is integrated only in V4, and this information is then inherited by IT via feed-forward propagation. **b)** The class of “IT: integrated” proposals predict that top-down information is integrated directly in IT. This class includes proposals in which top-down information is integrated in both IT and V4 (red) as well as proposals in which top-down information is integrated exclusively in IT but is then fed-back to V4 (cyan).

**Figure 2.**
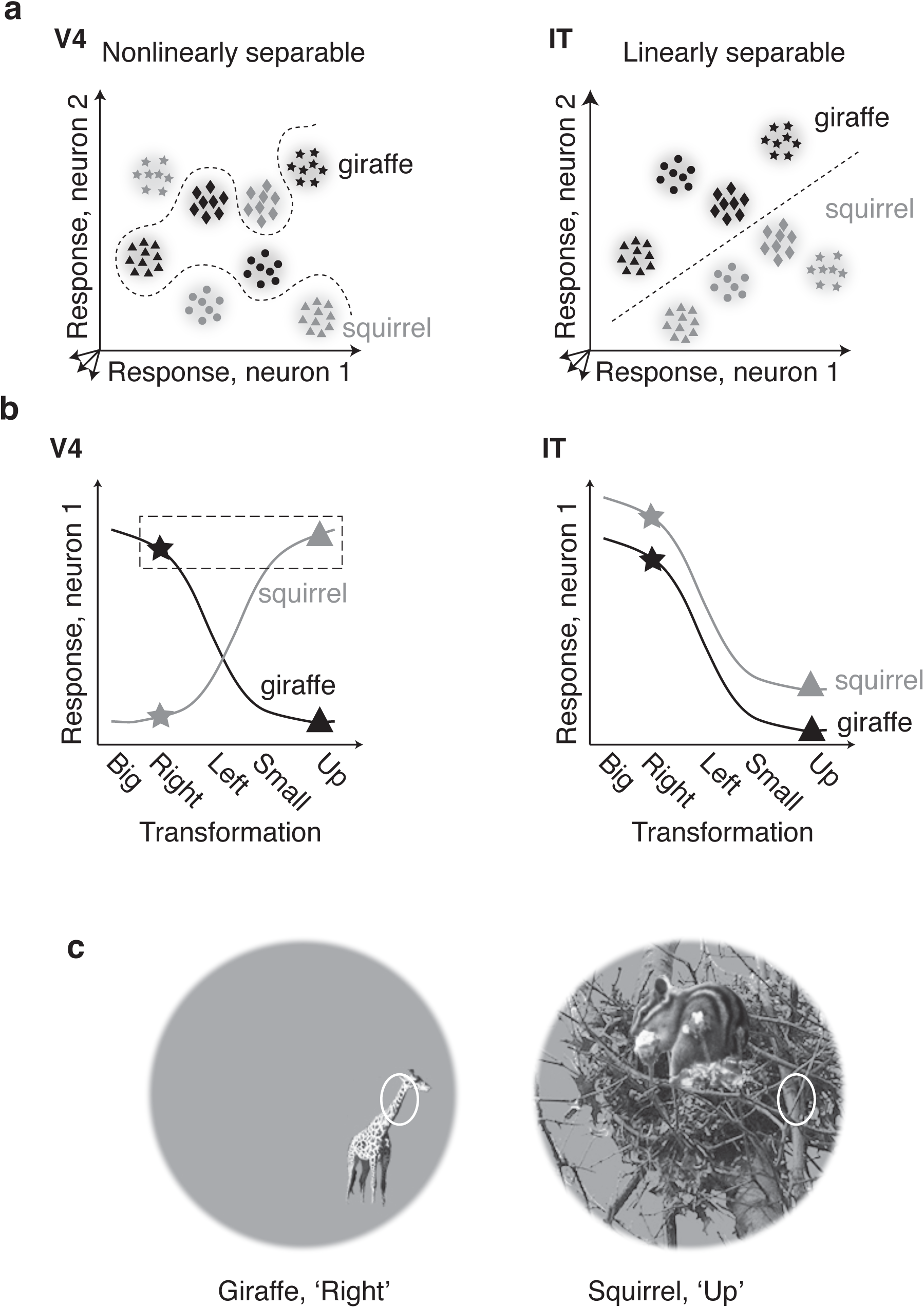
Differences between V4 and IT visual object representations. **a)** Cartoon depiction of the population representation of object identity in V4 and IT plotted as the response of two hypothetical neurons. In these plots, each point indicates the population response on a single trial, each cloud of points depicts the dispersion of responses to the same image across repeated trials, the shade of the points indicate the identity of the object contained in each image, and the shape of the points indicate the identity-preserving transformation at which each object appears. In V4, representations of object identity are nonlinearly separable, as indicated by the nonlinear boundary required to separate the two objects. In IT, representations of object identity are more linearly separable, as indicated by the linear boundary. **b)** The single-neuron property underlying V4 nonlinear and IT linear population representations is thought to be object tolerance, where increased IT linear separability (right) follows from the increased preservation of rank-order responses across identity-preserving transformations, and V4 nonlinear separability (left) follows from larger changes in rank-order preferences across different transformations. **c)** Illustration of how rank-order preferences for objects can change when receptive fields are small and objects are moved to a new position. In the left image, a region of giraffe object drives the neuron to fire; in the right image, the same neuron is driven by the background of an image containing a different object (squirrel).

The means by which the brain integrates top-down information in the ventral visual pathway may very well depend on the specific task and its computational demands. The experiments described here were targeted at challenging the hypothesis that top-down integration happens exclusively in the ventral visual pathway within or before V4 (Fig 1a) with a task that is seemingly better optimized for top-down integration in IT. Specifically, our experiments are designed to exploit differences in the visual representation of object identity between these two brain areas.

## MATERIALS AND METHODS

### Experimental Design

Experiments were performed on two adult male rhesus macaque monkeys (*Macaca mulatta)* with implanted head posts and recording chambers. All procedures were performed in accordance with the guidelines of the University of Pennsylvania Institutional Animal Care and Use Committee.

### Visual stimuli

Images were presented in a circular aperture with a radius of 5 degrees, centered at fixation (Fig 3a). Images contained objects presented at different positions, sizes and background contexts (Fig 3c). Sizes included: size-1x (∼1.2 degrees), size-1.5x (∼1.8 degrees) and size-2.25x (∼2.7 degrees). Positions included position-fixation (0 degrees horizontal and vertical), position-right (1.55 degrees to the right, 0.67 degrees below fixation), position-left (1.55 degrees to the left, 0.67 degrees below fixation) and position-up (0 degrees horizontal; 1 degree above fixation). Changes in size and position were combined to produce the following five transformations (Fig 3c): “Up” (position-up, size-1.5x); “Left” (position-left; size-1.5x); “Right” (position-right; size-1.5x); “Big” (position-fixation; size-2.25x); and “Small” (position-fixation; size-1x). In addition, a different natural image background was chosen for each of the transformations Up, Big and Small, whereas Left and Right were presented on a gray background (Fig 3c). The complete image set included 4 objects, each presented at the five transformations described above, for a total of 20 images. The rationale behind selecting these particular transformations was to make the task of object identification (invariant to identity-preserving transformation) challenging for both V4 and IT, based on results from our previous work (Rust and DiCarlo 2010).

**Figure 3.**
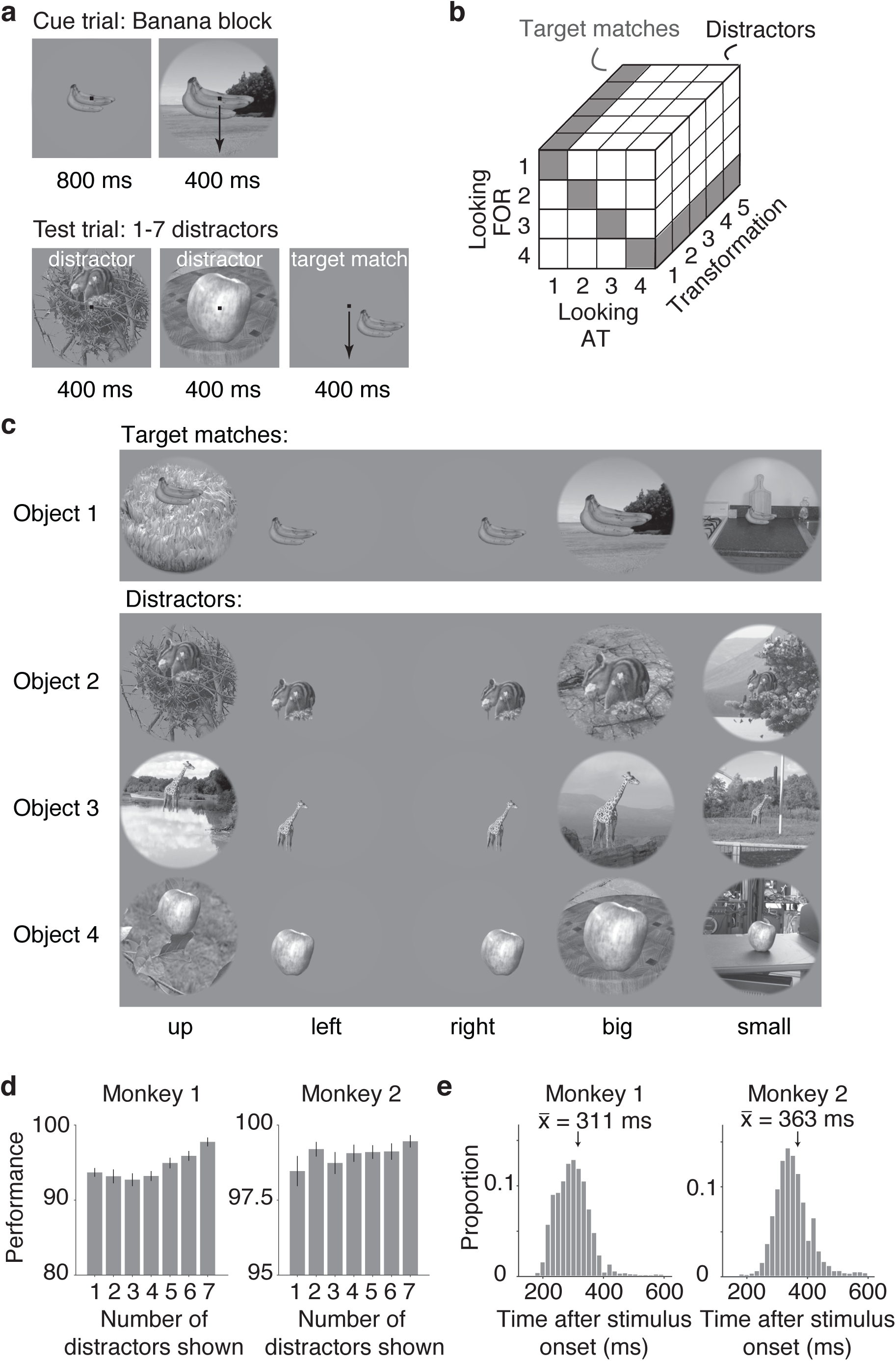
The invariant delayed-match-to-sample task (IDMS). **a)** Monkeys initiated trials by fixating on a small dot. Each block (∼3 minutes in duration) began with a cue trial which indicated the target object. On subsequent trials, a random number (1-7) of distractors were presented, and on most trials, this was followed by the target match. Monkeys were required to maintain fixation throughout the distractors and make a saccade to a response dot within a window 75 - 600 ms following the onset of the target match to receive a reward. In cases where the target match was presented for 400 ms and the monkey had still not broken fixation, a distractor stimulus was immediately presented. **b)** A schematic of the full experimental design, which included 80 conditions: looking “at” each of 4 objects, each presented at 5 identity-preserving transformations (for 20 images in total), viewed in the context of looking “for” each object as a target. In this design, target matches (gray) fall along the diagonal of each “looking at” / “looking for” transformation slice whereas distractors (white) fall off the diagonal. **c)** Images used in the task: 4 objects were presented at each of 5 identity-preserving transformations (“up”, “left”, “right”, “big”, “small”), for 20 images in total. In any given block, 5 of the images were presented as target matches and 15 were distractors. **d)** Percent correct for each monkey, calculated based on both misses and false alarms (but disregarding fixation breaks), shown as a function of the number of distractors preceding the target match. Error bars indicate standard error across experimental sessions. **e)** Histograms of reaction times during correct trials (ms after stimulus onset), with means labeled.

As described below, our experiment included two types of trials, “Cue trials” and “Test trials”. The set of 20 images described above were the only images presented on Test trials. Cue trials also included a version of each object (size-1.5x) presented at position-fixation on a gray background.

#### The invariant delayed-match-to-sample (IDMS) task

Monkey behavioral training and testing utilized standard operant conditioning, head stabilization and infrared video eye tracking. Custom software (https://mworks.github.io/) was used to present stimuli on an LCD monitor with an 85 Hz refresh rate.

The monkeys performed an invariant delayed-match-to-sample task (Fig 3). As an overview, the task required the monkeys to make a saccade when a target object appeared within a sequence of distractor images (Fig 3a). Objects were presented at differing positions, sizes and background contexts as described above and shown in Figure 3c. Stimuli consisted of a fixed set of 20 images that included 4 objects, each presented at 5 different identity-preserving transformations (Fig 3b). Each short block (∼3 min) was run with a fixed target object before another target was pseudorandomly selected. Our design included two types of trials: cue trials and test trials (Fig 3a). Only test trials were analyzed for this report.

A trial began when the monkey fixated on a red dot (0.15°) in the center of a gray screen, within a square window of ±1.5°. Fixation was followed by a 250 ms delay before a stimulus appeared. Cue trials, which indicated the current target object, were presented at the beginning of each short block or after three subsequent error trials. To minimize confusion, cue trials were designed to be distinct from test trials and began with the presentation of an image of each object that was distinct from the images used on test trials (a large version of the object presented at the center of gaze on a gray background; Fig 3a). Test trials began with a distractor image, and neural responses to the first distractor were discarded to minimize non-stationarities such as stimulus onset effects. During the IDMS task, all images were presented at the center of gaze, in a circular aperture that blended into a gray background (Fig 3c).

In each block, 5 images were presented as target matches and the other 15 as distractors. Distractor images were drawn randomly without replacement until each distractor was presented once on a correct trial, and the images were then re-randomized. On most test trials, a target match followed the presentation of a random number of 1-6 distractors (Fig 3a). On a small fraction of trials, 7 distractors were shown, and the monkeys were rewarded for fixating through all distractors. Each image was presented for 400 ms (or until the monkeys’ eyes left the fixation window) and was immediately followed by the presentation of the next stimulus. Monkeys were rewarded for making a saccade to a response target within a window of 75 – 600 ms after the target match onset. In monkey 1, the response target was positioned 10 degrees below fixation; in monkey 2 it was 10 degrees above fixation. If the monkeys had not yet moved their eyes after 400 ms following target onset, a distractor stimulus was immediately presented. A trial was classified as a ‘false alarm’ if the eyes left the fixation window via the top (monkey 1) or bottom (monkey 2) outside the allowable correct response period and travelled more than 0.5 degrees. In contrast, all other instances in which the eyes left the fixation window during the presentation of distractors were characterized as fixation breaks. A trial was classified as a ‘miss’ when the monkey continued fixating beyond 600 ms following the onset of the target match. Within each block, 4 repeated presentations of each of the 20 images were collected, and a new target object was then pseudorandomly selected. Following the presentation of all 4 objects as targets, the targets were re-randomized. At least 10 repeats of each condition were collected on correct trials. When more than 10 repeats were collected, the first 10 were used for analysis. Overall, monkeys performed this task with high accuracy. Disregarding fixation breaks (monkey 1: 8% of trials, monkey 2: 11% of trials), percent correct on the remaining trials was: monkey 1: 94% correct, 2% false alarms, and 4% misses; monkey 2: 98% correct, ∼1% false alarms, and ∼1% misses. Behavioral performance was comparable for the sessions corresponding to recordings from the two areas (V4 percent correct overall = 96.5%; IT percent correct overall = 91.4%).

V4 receptive fields at and near the center of gaze are small: on average they have radii of 0.56 degrees at the fovea, extending to radii of 1.4 at an eccentricity of 2.5 degrees (Desimone and Schein 1987; Gattass et al. 1988). We thus took considerable care to ensure that that the images were approximately placed in the same region of these receptive fields across repeated trials. In the second monkey, adequate fixational control could not be achieved through training. We thus applied a procedure in which we shifted each image at stimulus onset 25% toward the center of gaze (e.g. if the eyes were displaced 0.5 degrees to the left, the image was repositioned 0.125 degrees to the left and thus 0.375 degrees from fixation). Image position then remained fixed until the onset of the next stimulus.

### Neural recording

The activity of neurons in V4 and IT was recorded via a single recording chamber for each brain area in each monkey. In both monkeys, chamber implantation and recording in IT preceded V4, and the IT recording chamber was implanted on the right hemisphere whereas the V4 recording chamber was implanted on the left hemisphere. While IT receptive fields span the vertical meridian, thus allowing us to access the visual representation of both sides with a single chamber, V4 receptive fields are confined to the contralateral hemifield. To simulate V4 coverage of the ipsilateral visual field, on roughly half of the V4 recording sessions, (n = 7/15 sessions in Monkey 1, n = 11/20 sessions in Monkey 2), we presented the images reflected across the vertical axis. We then treated all V4 units recorded during these sessions as if they were in the left hemisphere (and thus as receptive fields that were located in the right visual field).

Chamber placement for the IT chambers was guided by anatomical magnetic resonance images in both monkeys, and in one monkey, Brainsight neuronavigation (https://www.rogue-research.com/). Both V4 chambers were guided by Brainsight neuronavigation. The region of IT recorded was located on the ventral surface of the brain, over an area that spanned 4 mm lateral to the anterior middle temporal sulcus and 15-19 mm anterior to the ear canals. Both V4 chambers were centered 1 mm posterior to the ear canals and 29 mm lateral to the midline, positioned at a 30 degree angle. V4 recording sites were confirmed by a combination of receptive field location and position in the chamber, corresponding to results reported previously (Gattass et al. 1988). Specifically, we recorded from units within and around the inferior occipital sulcus, between the lunate sulcus and superior temporal sulcus. V4 units in lower visual field were confirmed as having receptive field centers that traversed from the vertical to horizontal meridian as recordings shifted from posterior to anterior. As expected, V4 units in the fovea and near the upper visual field were found lateral to those in the lower visual field, and had receptive field centers that traversed from the horizontal meridian to the vertical meridian as recordings traversed medial to lateral and increased in depth.

Neural activity was recorded with 24-channel U-probes and V-probes (Plexon, Inc) with linearly arranged recording sites spaced with 100 mm intervals. Continuous, wideband neural signals were amplified, digitized at 40 kHz and stored using the OmniPlex Data Acquisition System (Plexon, Inc.). Spike sorting was done manually offline (Plexon Offline Sorter). At least one candidate unit was identified on each recording channel, and 2-3 units were occasionally identified on the same channel. Spike sorting was performed blind to any experimental conditions to avoid bias. A multi-channel recording session was included in the analysis if the animal performed the task until the completion of at least 10 correct trials per stimulus condition, there was no external noise source confounding the detection of spike waveforms, and the session included a threshold number of task-modulated units (>4 on 24 channels). The sample size for IT (number of units recorded) was chosen to approximately match our previous work (Pagan et al. 2013; Pagan and Rust 2014a). The sample size for V4 was selected to be 3-fold that number, to match the ratio between numbers of units estimated in V4 as compared to IT (DiCarlo et al. 2012).

For many of the analyses presented in this paper, we measured neural responses by counting spikes in a window that began 40 ms after stimulus onset in V4 and 80 ms after stimulus onset in IT. We counted spikes in a 170 ms window in both areas, such that the spike counting windows were of equal length. Counting windows always preceded the monkeys’ reaction times.

On 7.7% of all correct target match presentations, the monkeys had reaction times faster than 250 ms, and those instances were excluded from analysis to ensure that spikes in both V4 and IT were only counted during periods of fixation.

In IT, we recorded neural responses across 20 experimental sessions (Monkey 1: 10 sessions, and Monkey 2: 10 sessions). In V4, we recorded neural responses across 35 experimental sessions (Monkey 1: 15 sessions, and Monkey 2: 20 sessions). When combining the units recorded across sessions into a larger pseudopopulation, we began by screening for units that met three criteria. First, units needed to be modulated by our task, as quantified by a one-way ANOVA applied to our neural responses (80 conditions * 10 repeats, p < 0.01). Second, units needed to pass a loose criterion on recording stability, as quantified by calculating the variance-to-mean ratio (Fano factor) for each unit, computed by fitting the relationship between the mean and variance of spike count across the 80 conditions (Fano factor < 5). Finally, units needed to pass a loose criterion on unit recording isolation, quantified by calculating the signal-to-noise ratio (SNR) of the waveform as the difference between the maximum and minimum points of the average waveform, divided by twice the standard deviation across the differences between each waveform and the mean waveform (SNR > 2). In IT, this yielded a pseudopopulation of 204 units (of 563 possible units), including 108 units from monkey 1 and 96 units from monkey 2. In V4, this yielded a pseudopopulation of 650 units (of 970 possible units), including 382 units from monkey 1 and 268 units from monkey 2.

### V 4 receptive field mapping

To measure the location and extent of V4 receptive fields, bars were presented for 500 ms, one per trial, centered on a 5 × 5 invisible grid. Bar orientation, length, and width as well as the grid center and extent were adjusted for each recording session based on preliminary hand mapping. On each trial, the monkey was required to maintain fixation on a small response dot (0.125°) to receive a reward. The responses to at least five repeats were collected at each position for each recording session. Only those units that produced clear visually evoked responses at a minimum of one position were considered for receptive field position analysis. The center of the receptive field was estimated by the maximum of the response across the 5×5 grid of oriented bar stimuli and confirmed by visual inspection.

#### Quantifying single-unit modulations

To quantify the degree to which individual V4 and IT units were modulated by task-relevant variables (Figs 6, 10, 11), such as changes in visual and target identity, we applied a bias-corrected, ANOVA-like procedure described in detail by (Pagan and Rust 2014b) and summarized here. As an overview, this procedure is designed to parse each unit’s total response variance into variance that can be attributed to each type of experimental parameter as well as variance that can be attributed to trial variability. Total variance is computed across the spike count responses for each unit across 16 conditions (4 images * 4 targets separately for each transformation) and 10 trials. Variances are then transformed into measures of spike count modulation (in the units of standard deviation around each unit’s grand mean spike count) via a procedure that includes bias correction for over-estimates in modulation due to noise.

To capture all types of modulation with intuitive groupings, the procedure begins by developing an orthonormal basis of 16 vectors. The number of basis vectors for each type of modulation is imposed by the experimental design. In particular, this basis ***b*** includes vectors ***b***_*i*_ that reflect 1) the grand mean spike count across all conditions, 2) whether the object in view is a target or a distractor (‘target match’), 3) visual image identity (‘visual’), 4) target object identity (‘target identity’), and 5) nonlinear interactions between target and object identity not captured by target match modulation (‘residual’). The initially designed set of vectors is then converted into an orthonormal basis via a Gram-Schmidt orthogonalization process.

The resulting basis spans the space of all possible responses for our task. Consequently, we can re-express each trial-averaged vector of spike count responses to the 16 experimental conditions for each transformation, ***R***, as a weighted sum of these basis vectors. The weight corresponding to a basis vector for each unit reflect modulation of that unit’s responses by that experimental parameter. To quantify the amounts of each type of modulation reflected by each unit, we began by computing the squared projection of each basis vector ***b***_*i*_ and ***R***. To correct for bias caused by over-estimates in modulation due to noise, an analytical bias correction, described and verified in (Pagan and Rust 2014b), was then subtracted from this value. The squared weight for each basis vector ***b***_*i*_ is calculated as:

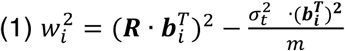

where 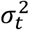 indicates the trial variance, averaged across conditions (n=16), and m indicates the number of trials (m=10). If more than one dimension existed for a type of modulation, we summed values of the same type (eq. 2). Next, we applied a normalization factor (1/(n-1)) where n=16) to convert these summed values into variances. As a final step, we computed the square root of these quantities to convert them into modulation measures that reflected the number of spike count standard deviations around each unit’s grand mean spike count. Modulation for each parameter type X was thus computed as:

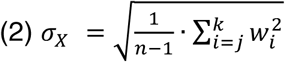

for the weights *w*_*j*_ through *w*_*k*_ corresponding to basis vectors ***b***_*j*_ through ***b***_*k*_ for that parameter type, where the number of basis vectors corresponding to each parameter type were: target match = 1; visual = 3; target identity = 3; residual = 8.

When estimating modulation for individual units, (Fig 6), the bias-corrected squared values were rectified for each unit before taking the square root. When estimating modulation population means (Figs 10b-e, 11), the bias-corrected squared values were averaged across units before taking the square root. When estimating modulation population means within the broader 170 ms bins for V4 and IT respectively (bar graphs shown in Figs 10b-e, 11 right), the modulations shown are roughly equal to the sum (or integral) of the modulations in the 50 ms sliding bins (Figs 10b-e, 11, left) summed across the 40-20 ms or 80-250 ms bins in V4 and IT respectively. However, because we computed the modulations separately in the different bin sizes before bias-correcting, averaging across units, and taking the square root, this relationship is not exactly equivalent to an integral. Because these measures were not normally distributed, standard error about the mean was computed via a bootstrap procedure. On each iteration of the bootstrap (across 1000 iterations), we randomly sampled values from the modulation values for each unit in the population, with replacement. Standard error was computed as the standard deviation across the means of these resampled populations.

### Population performance: Visual object invariance

To determine performance of the V4 and IT populations at classifying visual object identity (Fig 7), we computed 4-way object discrimination performance. As an overview, we formulated the problem as four one-versus-rest linear classifications, and then took the maximum of these classifications as the population’s decision (Hung et al. 2005). Here we begin by describing the general form of linear classifier that we used, a Fisher Linear Discriminant (FLD), and we then describe the training and testing scheme for measuring cross-validated performance.

The general form of a linear decoding axis is:

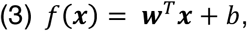

where **w** is an N-dimensional vector containing the linear weights applied to each of N units, and b is a scalar value. We fit these parameters using an FLD, where the vector of linear weights was calculated as:

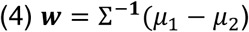

and b was calculated as:

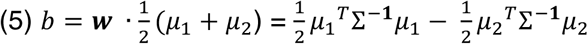

Here *μ*_1_ *and μ*_2_ are the means of two classes (e.g. two object classes, respectively) and the mean covariance matrix is calculated as:

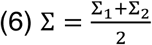

where Σ_1_ and Σ_2_ are the regularized covariance matrices of the two classes. These covariance matrices were computed using a regularized estimate equal to a linear combination of the sample covariance and the identity matrix *I* (Pagan and Rust 2014a):

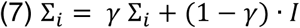

We determined *γ* by exploring a range of values from 0.01 to 0.99, and we selected the value that maximized average performance across all iterations, measured with the cross-validation “regularization” trials set aside for this purpose (see below). We then computed performance for that value of *γ* with separately measured “test” trials, to ensure a fully cross-validated measure. Because this calculation of the FLD parameters incorporates the off-diagonal terms of the covariance matrix, FLD weights are optimized for both the information conveyed by individual units as well as their pairwise interactions.

To classify which of four objects was in view, we used a standard “one-versus-rest” classification scheme. Specifically, one linear classifier was determined for each object based on the training data. To determine the population decision about which object was presented, a response vector **x,** corresponding to the population response to one of the four objects, was then applied to each of the classifiers, and the classifier with the largest output (the classifier with the largest positive *f*(**x)**) was taken as the population decision. To train the classifiers, we used an iterative resampling procedure. On each iteration of the resampling, we randomly shuffled the trials for each condition and for each unit, and (for numbers of units less than the full population size) randomly selected units. On each iteration, 8 (of the 10 total) trials from each condition were used for training the decoder, 1 trial from each condition was used to determine a value for regularization, and 1 trial from each condition was used for cross-validated measurement of performance.

We compared classifier performance for the “reference” cases (when cross-validated test trials were selected from the same transformation used to train the classifier; Fig 7a-c, black) versus the “generalization” cases (when test trials were selected from transformations different than the one used for training, Fig 7a-c, cyan). To summarize the results for a given transformation, reference and generalization performance was compared for the same test data: e.g. in the case of the transformation “Up”, reference performance was computed by training and cross-validated testing on “Up” and generalization performance was computed as the average of training on all other transformations and testing on “Up”.

To ensure that visual classification performance was not biased by the target match signal, we computed performance for targets and distractors separately and averaged their results. Specifically, we computed visual classification performance for the four objects presented as target matches or for different combinations of the four objects presented as distractors. Each set of 4 distractors was selected to span all possible combinations of mismatched object and target identities (e.g. objects 1, 2, 3, 4 paired with targets 4, 3, 2, 1), of which there are 9 possible sets. As a final measure of visual classification performance, we then averaged across 10 performance values (1 target match and 9 distractor combinations) as well as, when relevant, multiple transformations. One performance value was computed on each iteration of the resampling procedure, and mean and standard error of performance was computed as the mean and standard deviation of performance across 1000 resampling iterations. Standard error thus reflected the variability due to the specific trials assigned to training and testing and, for populations smaller than the full size, the specific units chosen. Finally, generalization capacity was computed on each resampling iteration by taking the ratio of the chance-subtracted reference performance and the chance-subtracted generalization performance (where chance = 25%).

### Population performance: Target match information

To determine the ability of the V4 and IT populations to classify target matches versus distractors (Fig 8), we applied two types of decoders: a linear classifier (an FLD, described above) and a Maximum Likelihood decoder (a decoder that can classify based on linear as well as nonlinearly formatted target match information). Both decoders were cross-validated with the same resampling procedure. On each iteration of the resampling, we randomly shuffled the trials for each condition and for each unit, and (for numbers of units less than the full population size) randomly selected units (with the exception of Fig 8b, cyan, where we selected the ‘best’ units, as described below). On each iteration, 8 (of the 10 total) trials from each condition were used for training the decoder, 1 trial from each condition was used to determine a value for regularization of the FLD linear classifier (see below) and 1 trial from each condition was used for a cross-validated measurement of performance.

To circumvent issues related to the format of visual information, classifier analyses were performed per transformation (for the 4 of the 5 transformations used, see Fig 6; “Big”, “Up”, “Left” and “Small”). The data for each transformation consisted of 16 conditions (4 visual objects viewed under 4 different target contexts). To ensure that decoder performance relied only on target match information and not on other factors, such as differences in the numbers of each class, each classification was computed for 4 target matches versus 4 (of 12 possible) distractors. Each set of 4 distractors was selected to span all possible combinations of mismatched object and target identities (e.g. objects 1, 2, 3, 4 paired with targets 4, 3, 2, 1), of which there are 9 possible sets. Performance was computed on each resampling iteration by averaging the binary performance outcomes across the 9 possible sets of target matches and distractors, each which contained 8 cross-validated test trials, and across the four transformations used. For both types of classifiers, mean and standard error of performance was computed as the mean and standard deviation of performance across 1000 resampling iterations. Standard error thus reflected the variability due to the specific trials assigned to training and testing and, for populations smaller than the full size, the specific units chosen.

To compute linear classifier performance (Fig 8b), we used a 2-way Fisher Linear Discriminant, described as in the general form above. In this case, the classes described in eqs. 4-6 correspond to target matches and distractors. To compute neural population performance, we began by computing the dot product of the test data and the linear weights **w,** adjusted by *b* (Eq. 3). Each test trial was then assigned to one class, and proportion correct was then computed as the fraction of test trials that were correctly assigned, according to their true labels. To compute linear classifier performance for the best V4 units (Fig 8b, cyan), we ranked units by their d’ based on the training data and sub-selected top-ranked units to measure cross-validated performance. Unit d’ was computed as:

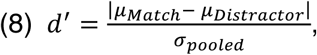

where *μ*_*Match*_ and *μ*_*Distractor*_ correspond to the mean response across the set of target match and distractors, 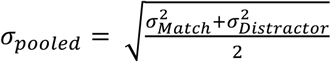 and *σ*_*Match*_ and *σ*_*Distractor*_ correspond to the standard deviation of responses across the set of target matches and distractors, respectively.

As a measure of total target match information (Fig 8d; combined linear and nonlinear), we implemented a maximum likelihood decoder (Pagan et al. 2013; Pagan et al. 2016). We began by using the set of training trials to compute the average response r_uc_ of each unit u to each of the 2 conditions c (target matches versus distractors). We then computed the likelihood that a test response k was generated from a particular condition as a Poisson-distributed variable:

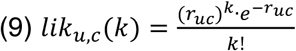

The likelihood that a population response vector was generated in response to each condition was then computed as the product of the likelihoods of the individual units. We assigned the population response to the category with the maximum likelihood, and we computed performance as the fraction of trials in which the classification was correct based on the true labels of the test data.

### Statistical analysis

Because our measures were not normally distributed, we computed P values via resampling procedures. When comparing the magnitudes of single unit modulation values between V4 and IT (Figs 6, 10b-e, 11), a bootstrap procedure was applied in which values were randomly sampled from the values for each unit, with replacement, across many iterations. We calculated *P* values as the fraction of resampling iterations on which the difference was flipped in sign relative to the actual difference between the means of the full data set (for example, if the mean of visual modulation in V4 was larger than the mean of visual modulation in IT, the fraction of iterations in which the mean of visual modulation in IT was larger than the mean of visual modulation in V4).

When comparing generalization capacity between the V4 and IT populations (Fig 7d), we began by computing generalization capacity for each of 1000 resampling iterations of the reference and generalization classifiers. We calculated *P* values as the fraction of resampling iterations on which the difference was flipped in sign relative to the actual difference between the means of the full data set (for example, if the mean of generalization capacity in IT was larger than the mean of generalization capacity in V4, the fraction of iterations in which the mean of generalization capacity in V4 was larger than the mean of generalization capacity in IT).

When comparing population decoding measures (Figs 7a-c, 8b, & 8d), 1000 iterations of cross-validated population performance were computed, and *P* values were calculated as the fraction of classifier iterations on which the difference was flipped in sign relative to the actual difference between the means across classifier iterations (for example, if the mean of decoding measure 1 was larger than the mean of decoding measure 2, the fraction of iterations in which the mean of measure 2 was larger than the mean of measure 1). When evaluating whether a population decoding measure was different from chance (Fig 8), P values were calculated as the fraction of classifier iterations on which performance was greater than chance performance (50%).

## RESULTS

Understanding the path by which top-down information arrives in the ventral visual pathway during visual object search is challenging, due to several factors. First, receptive fields at different stages of the pathway have markedly different sizes and it’s unclear how to best compare them. Second, during tasks that approximate real-world object search, the responses of individual V4 and IT neurons operate in a low spike count regime in which the amounts and types of signals within individual units are difficult to measure (Roth and Rust 2018a; Roth and Rust 2018b). This noisy, low spike count regime is a combined consequence of spike count windows that are short, as implied by reaction times that are fast (∼250 ms), and individual stimuli that drive only a subset of neurons robustly (Roth and Rust 2018b). These challenges have traditionally been addressed using approaches that seek to reduce the noise in a manner that implicitly makes unrealistic assumptions about the brain. For example, tailoring stimuli to fit within the small sizes of V4 receptive fields and/or aligning stimuli with the peak of each neuron’s preferences disregards the contributions of the receptive field surround and/or neurons that are activated along their curve flank. Similarly, counting spikes in long windows that exceed natural reaction times assumes that neural responses are stationary, whereas their responses are not (e.g. Pagan and Rust 2014b). Here we apply a complementary approach that involves studying V4 and IT in a manner analogous to the way that the brain addresses the challenge of noisy individual neuron responses during real-world object search: by combining the noisy responses of individual neurons across a neural population (i.e. via weighted population decoding schemes).

As an overview of our experiments, we trained two monkeys to perform an “invariant delayed-match-to-sample” (IDMS) task that required them to report when target objects appeared. On any given trial of the IDMS task, monkeys were instructed about the object that they were searching for, but they did not know the specific position, size and background context with which it would appear. The task was designed to exploit well-established differences in the nature of visual object representations between V4 and IT, where both brain areas have been demonstrated to reflect similar total amounts of information for object identification tasks, but in V4 this information is more implicitly formatted (i.e. nonlinearly separable; Fig 2a, left) whereas in IT this information is more explicitly formatted (i.e. more accessible to a linear population read-out; Fig 2a right; reviewed by DiCarlo et al. 2012). At the level of individual neurons, the response property supporting IT population linear separability is thought to be object “tolerance”, or the preservation of rank-order tuning for objects across identity-preserving transformations (Ito et al. 1995; Li et al. 2009). Object tolerance is most intuitively described for changes in object position and spatially large IT receptive fields that have the same tuning for object identity across all regions of their receptive field (Fig 2b, right). In contrast, V4 receptive fields are smaller, and are stimulated by different regions of the image when an object moves to a new position (Fig 2c), and this can lead to a change in the rank-order tuning for objects at different positions (Fig 2b, left).

In the context of the IDMS task where monkeys were searching for an object that could appear at different transformations, the visual information required to solve the task (i.e. identity of the object in view) is expected to exist in both V4 and IT, but this information is expected to be formatted in a more linear manner in IT (Fig 2a). In question is whether this difference in information format between V4 and IT coincides with the preferential integration of top-down information in IT (Fig 1b). Equivalently, in question is whether during the IDMS task the brain somehow manages to integrate invariant information about object identity in V4 where receptive fields are small – which it may in fact be able to do, given the spatially global top-down modulations documented in V4 feature-based attention experiments (Maunsell and Treue 2006; Cohen and Maunsell 2011) - or whether it preferentially targets IT, where receptive fields are larger.

The IT data presented here were also included in two earlier publications (Roth and Rust 2018a; Roth and Rust 2018b). There, we reported that during IDMS, neural signals in IT reflected behavioral confusions on the trials in which the monkeys made errors, and IT target match signals were configured in a manner that minimized their interference with IT visual representations. The focus of the current report is a determination of how these signals arrive in IT via a systematic comparison between IT and its input brain area, V4; the V4 data have not been published previously.

### The invariant delayed-match-to-sample task (IDMS)

Monkeys performed an invariant delayed-match-to-sample (IDMS) task in short blocks of trials (∼3 minutes on average) with a fixed target object. Each block began with a cue trial that indicated the target for that block (Fig 3a, ‘Cue Trial’). The remainder of the block was comprised primarily of test trials (Fig 3a, ‘Test trial’). Test trials began with the presentation of a distractor and on most trials this was followed by 0-5 additional distractors (for a total of 1-6 distractor images) and then an image containing the target match. The monkeys’ task required them to maintain fixation during the presentation of distractors and make a saccade in response to the appearance of a target match to receive a juice reward. To minimize the possibility that monkeys would predict the target match, on a small fraction of the trials the target match did not appear and the monkeys were rewarded for maintaining fixation through 7 distractors. Unlike other classic DMS tasks (Eskandar et al. 1992; Chelazzi et al. 1993; Leuschow et al. 1994; Miller and Desimone 1994; Pagan et al. 2013) our experimental design does not incorporate a cue at the beginning of each test trial, to better mimic real-world object search, where target matches are not repeats of the same image presented shortly before. One benefit of this task design is that it better isolates target match modulation from other types of stimulus repetition effects, including repetition suppression (Miller and Desimone 1994). This distinction is important for this study, as our goal was to systematically compare the magnitudes of top-down modulation in V4 and IT and repetition suppression is likely to be a largely feed-forward process.

Our experimental stimuli consisted of a fixed set of 20 images: 4 objects presented at each of 5 transformations (Fig 3c). These specific images were selected in order to make the task of classifying object identity challenging for the IT population and these specific transformations were selected based on findings from our previous work (Rust and DiCarlo 2010). In a given target block (e.g. a ‘banana block’), a subset of 5 of the images were target matches and the remaining 15 were distractors (Fig 3c). The full experimental design amounted to 20 images (4 objects presented at 5 identity-preserving transformations), all viewed in the context of each of the 4 objects as a target, resulting in 80 experimental conditions (Fig 3b). In this design, “target matches” fall along the diagonal of each “looking at” / “looking for” matrix slice (where a matrix “slice” corresponds to the conditions at one fixed transformation; Fig 3b, gray). For each of the 80 conditions, we collected at least 10 repeats on correct trials. Behavioral performance was high overall (Fig 3d). The monkeys’ mean reaction times (computed as the time their eyes left the fixation window relative to the target match stimulus onset) were 311 ms and 363 ms for monkey 1 and 2, respectively (Fig 3e).

To systematically compare the responses of V4 and IT during this task, we applied a population-based approach in which we fixed the images and their placement in the visual field across all the units that we studied, and we sampled from units whose receptive fields overlapped the stimuli. Specifically, we presented images at the center of gaze, with a diameter of 5 degrees. Neurons in IT typically have receptive fields that extend beyond 5 degrees and extend into all four quadrants (Fig 4a top; Op De Beeck and Vogels 2000). In contrast, V4 receptive fields are smaller, retinotopically organized, and confined to the contralateral hemifield (Fig 4a bottom; Desimone and Schein 1987; Gattass et al. 1988). To compare these two brain areas, we applied extensions of approaches developed in our earlier work in which we compared the responses of a set of randomly sampled IT units with a population of V4 units whose receptive fields tiled the image (Rust and DiCarlo 2010). This required sampling V4 units with receptive fields in both upper and lower visual fields (Fig 4b), which we achieved through recording at different positions within and around the inferior occipital sulcus. This also required measuring units with receptive fields on both sides of the vertical meridian, which we approximated by isolating our recordings to one hemisphere but reflecting the images along the vertical axis in approximately half the sessions.

**Figure 4.**
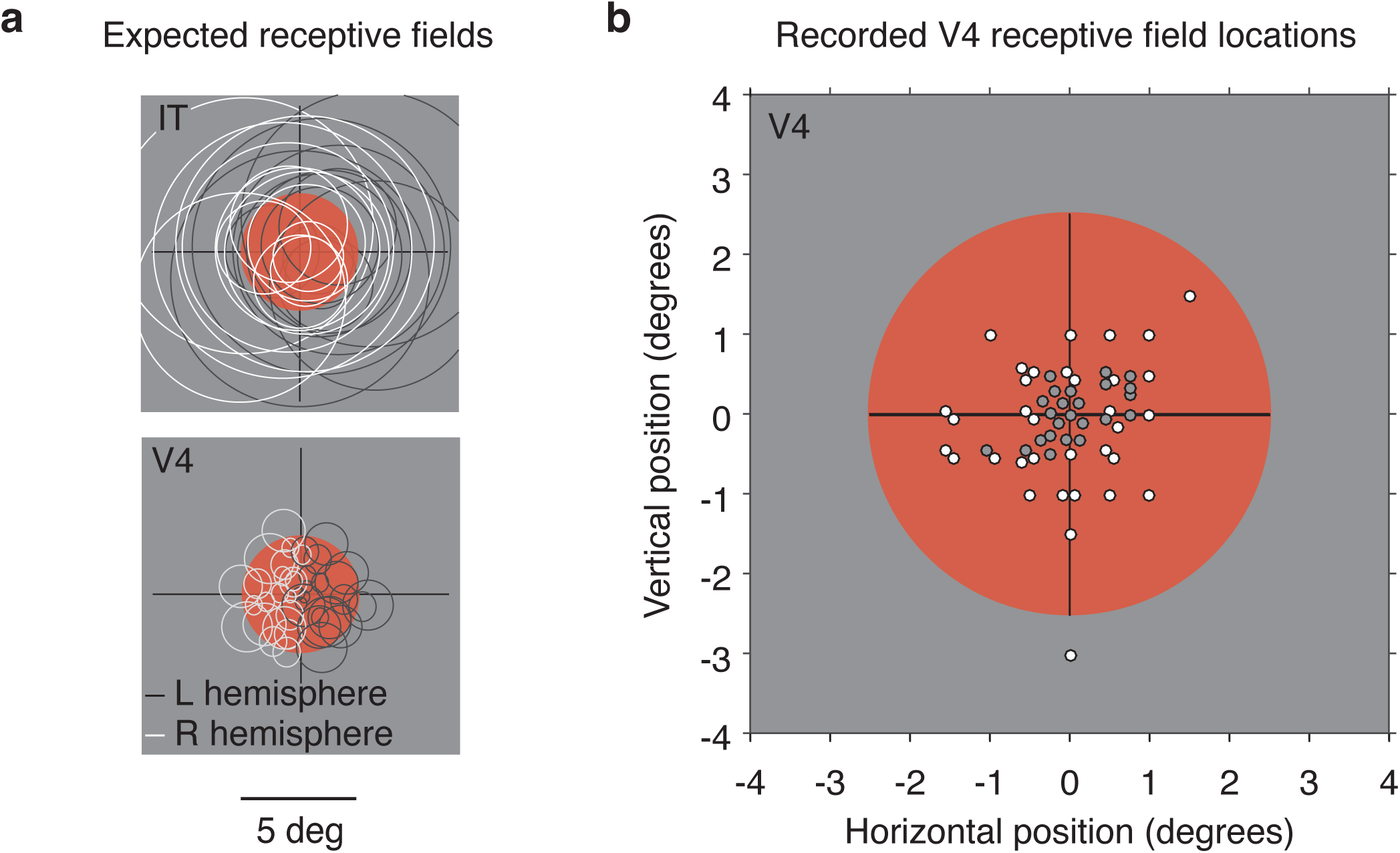
V4 and IT receptive field locations. Images were displayed at the center of gaze and were 5 degrees in diameter. Red circles indicate the location and size of the images. **a)** Schematic of expected receptive field locations and sizes for neurons in IT (top; Op De Beeck and Vogels 2000) and V4 (bottom; Desimone and Schein 1987; Gattass et al. 1988). **b)** We targeted V4 units with receptive fields that tiled the images. After approximate receptive field localization with hand mapping, receptive field locations were determined with oriented bar stimuli presented in a 5 × 5 grid of different positions (see Methods). Shown are the receptive field centers of a subset of recorded V4 units; one dot is shown for each unique receptive field location recorded. On approximately half of the sessions, images were reflected across the vertical axis, and for these sessions, the receptive field centers are plotted in the ipsilateral visual field. Monkey 1: gray; Monkey 2: white.

Because V4 receptive fields in the region of the field that we recorded are small, one issue of concern is the replicability of retinal image placement across trials. We quantified the stability of monkeys’ eye positions across repeated trials as the spatial deviation in retinal image placement across trials, measured relative to the mean position across trials, and we determined the proportion of eye positions that were within windows corresponding to V4 receptive field sizes at the range of eccentricities we recorded (Gattass et al. 1988). In monkey 1, 85% of eye positions fell within windows corresponding to the average RF sizes at the fovea (average foveal receptive field size = 0.56 degrees), and 97% of eye positions were within windows corresponding to RF sizes at an eccentricity of 2.5 degrees (average receptive field size at 2.5 degrees = 1.4 degrees). To achieve similar precision in Monkey 2, fixational control was improved by employing a procedure in which eye position was determined just before stimulus onset and the image was shifted at stimulus onset such that it was positioned closer to the center of gaze (see Methods). The image then remained in the same, fixed position for the duration of the image viewing period. The resulting retinal stability was comparable to monkey 1: on average, 95, and 99% of presentations occurred within windows with a radius of 0.56 and 1.4 degrees, respectively. These approaches were also effective in producing similar distributions of neural trial-by-trial variability between the two monkeys and between the two brain areas, as measured by the mean and standard deviation of the variance-to-mean ratio (Fano factor) across units (mean +/-std, Monkey 1: V4 = 1.61+/-0.65; IT = 1.62 +/-0.68; Monkey 2: V4 = 1.49 +/-0.54; IT = 1.28 +/-0.36).

As two monkeys performed this task, we recorded neural activity from small populations using 24-channel probes that were acutely lowered into V4 or IT before each session. With the rationale that V4 contains approximately 3-fold more units than IT near the fovea (DiCarlo et al. 2012), we aimed to collect 3-fold more units from V4. Following a screen for units based on their stability, isolation, and task modulation (see Methods), our data included 650 V4 units and 204 IT units (Monkey 1: 382 units in V4 and 108 in IT; Monkey 2: 268 units in V4 and 96 in IT). The data reported here were extracted from trials with correct responses. For most of our analyses, we counted spikes in equal length windows in V4 and IT but adjusted for the difference in latency between the two brain areas (170 ms, V4: 40-210 ms; IT: 80-250 ms following stimulus onset). These windows always preceded the monkeys’ reaction times and thus corresponded to periods of fixation. Distributions of peak, grand mean, and baseline firing rates computed for these spike count windows are shown in Figure 5. Consistent with a previous report (Rust and DiCarlo 2012), firing rates in V4 and IT were moderate, and counting spikes in the short windows implied by the monkeys’ fast reaction times only produced a few spikes per trial on average (mean firing rate was ∼1 spike/trial; V4: 1.34 spikes/trial; IT: 0.92 spikes/trial). In the face of these low spike counts, the statistical power in our analyses arises from combining the responses across many neurons using weighted decoding schemes.

**Figure 5.**
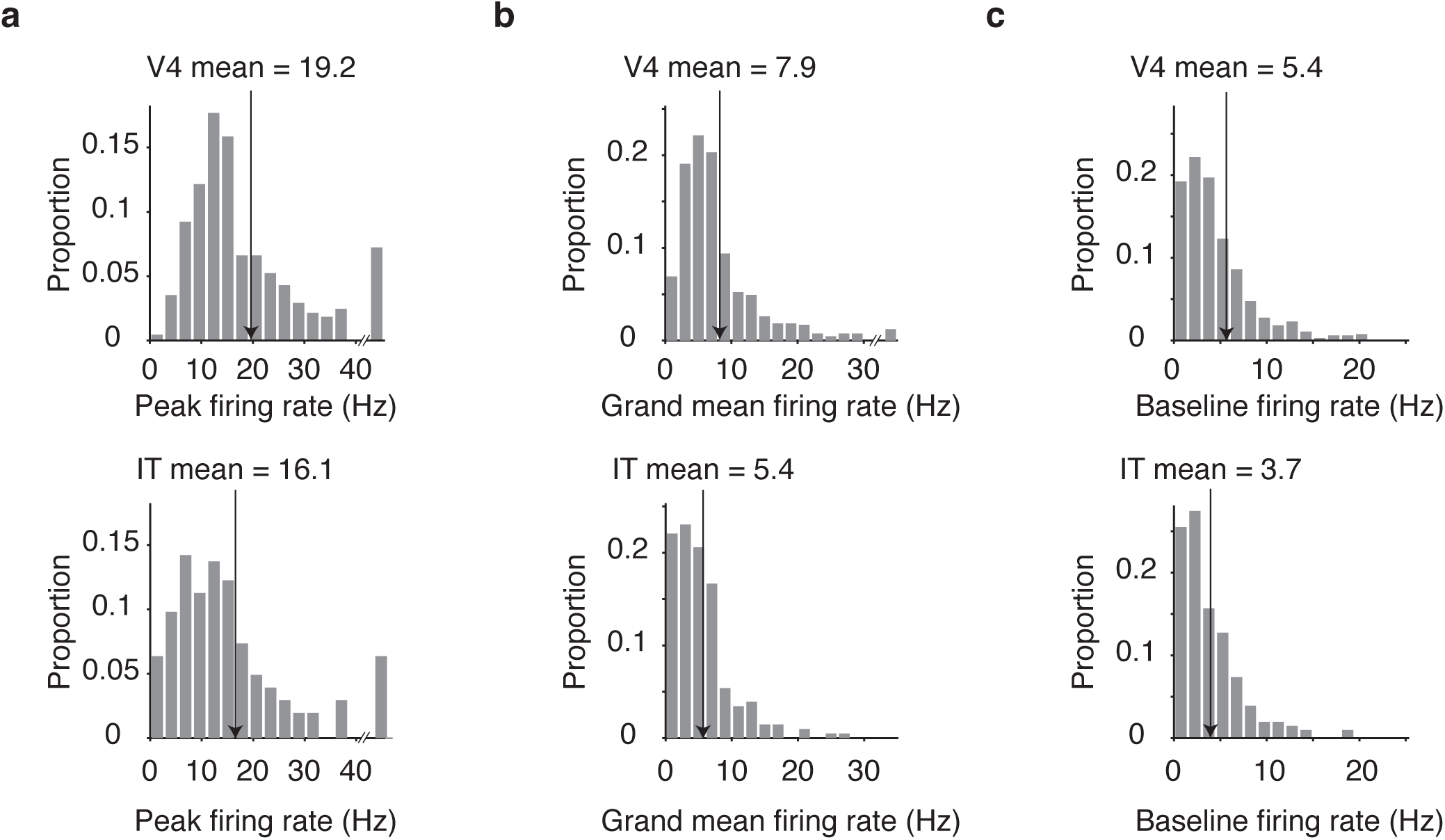
Firing rate distributions. Stimulus-evoked firing rates were computed as the average firing rate in a window 40-210 ms (V4) or 80–250 ms (IT) following stimulus onset whereas baseline firing rates were computed during the 170 ms period of fixation preceding stimulus onset. **a)** Maximum firing rates across the 80 stimulus conditions. The last bin includes units with maximum firing rates greater than 40 Hz. **b)** Grand mean firing rates across the 80 stimulus conditions. The last bin includes units with grand mean firing rates greater than 30 Hz.**c)** Baseline firing rates, computed in a 170 ms period of fixation before stimulus onset. In all panels, arrows indicate the means (V4: n = 650 units; IT: n = 204 units).

**Figure 6.**
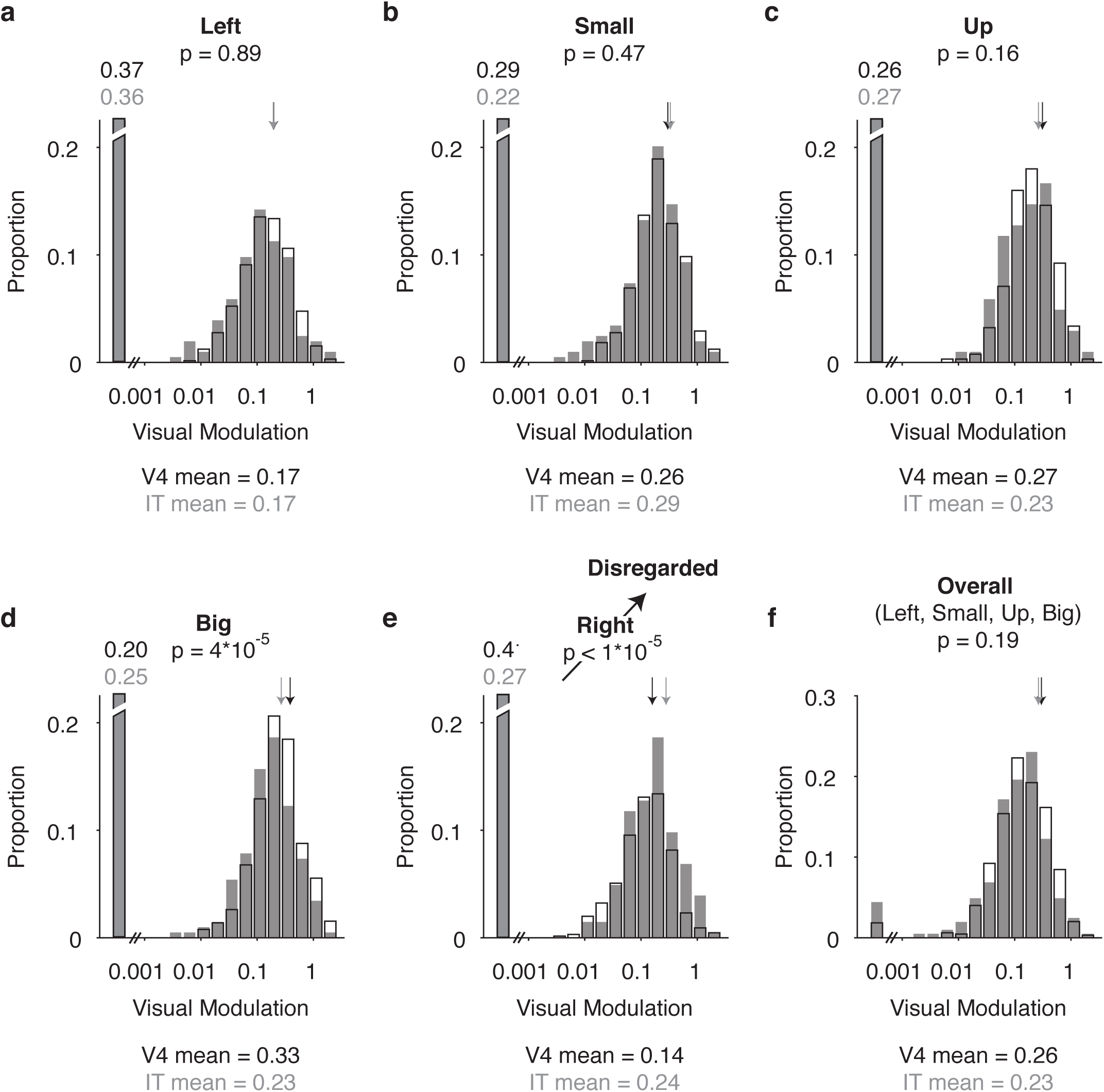
Comparison of visual modulation in V4 and IT. Shown are distributions of visual modulation magnitudes across units, parsed by transformation, for V4 (open bars, n = 650 units) and IT (gray, n = 204 units) and plotted on a log axis. Following a bias correction to remove the impact of trial variability, visual modulation was computed in units of standard deviation around each unit’s grand mean spike count. The first bin includes units with negligible visual modulation (modulation < 0.001) and the broken axis indicates that these bars should extend to the proportions labeled just above. Means of each distribution, including units with negligible visual modulation, are indicated by arrows and values are indicated at the bottom of each panel. The p-values at the top of each panel were computed via a bootstrap significance test evaluating the probability that differences in the means between V4 and IT can be attributed to chance. **a-e)** Distributions parsed by transformation. Visual modulation corresponding to the transformation ‘right’ was higher in IT as compared to V4, due to incomplete sampling of receptive fields at this location (Fig 4b), and was thus disregarded from further analyses. **f)** Distributions of visual modulation, averaged for each unit across the transformations ‘left’, ‘small’, ‘up’, and ‘big’.

**Figure 7.**
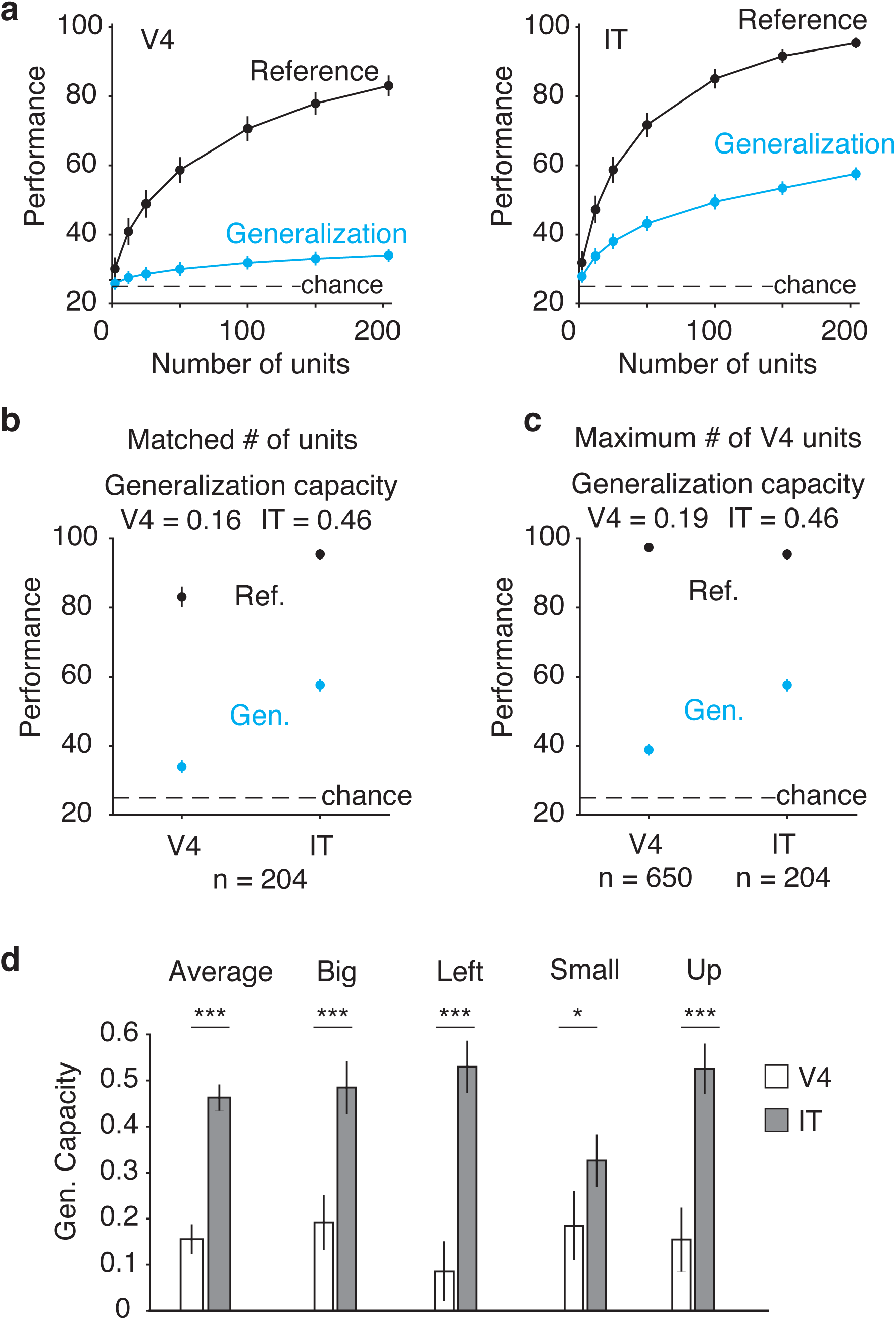
Comparison of visual object invariance across identity-preserving transformations in V4 versus IT. **a)** Performance of V4 and IT on a 4-way linear read-out of object identity, assessed either with cross-validated trials of the same transformation (“Reference”) or when asked to generalize to transformations not used for training (“Generalization”; see text). **b)** Reference and generalization performance for matched numbers of units (n = 204 for both V4 and IT populations), replotted from the endpoints in panel a. Generalization capacity was computed as the ratio of generalization over reference performance after subtracting the value expected by chance (where chance = 25%). **c)** Reference and generalization performance for the full recorded V4 population (n = 650 units) as compared to the full recorded IT population replotted from panel b (n = 204 units). **d)** Generalization capacity computed for matched numbers of units in V4 and IT (n = 204 units), applied to the average across transformations (as computed for panel b) and computed for each transformation separately. Single asterisks denote p < 0.05; double asterisks denote p < 0.01; triple asterisks denote p < 0.001. In all panels, error bars (standard error) reflect the variability that can be attributed to the specific subset of trials chosen for training and testing and, for subsets of units smaller than the full population, the specific subset of units chosen.

As an overview of our analyses, we begin by verifying that visual information in our recorded V4 and IT data is, as expected, i) matched in its total amounts between V4 and IT and ii) object identity is more linearly formatted in IT than in V4. Next, we evaluate the scenarios presented in Fig 1 by comparing the top-down information reflected in V4 and IT with a number of different population decoding schemes. Finally, we examine how top-down modulation is reflected in the responses of individual V4 and IT units, both as a function of time and as a function of visual tuning.

### Visual representations in V4 and IT during the IDMS task

#### Visual information is matched in V4 and IT during the IDMS task

When making systematic comparisons between V4 and IT, there are important factors to consider. For example, should the information contained in the V4 and IT populations be compared with equal numbers of units? Similarly, what are appropriate benchmarks for determining whether the samples recorded from each brain area are representative? As an example, imagine a scenario in which the information about whether an image is a target match or a distractor is reflected in both V4 and IT to the same degree, but the V4 neurons recorded in an experiment all have small, overlapping receptive fields confined to the same, small region of the visual field. In contrast, IT neurons, by virtual of their large receptive fields, would have access to much more of the visual field. From this data we might erroneously find that the magnitude of total target match information is larger in IT than V4 by way of non-representative sampling.

As a benchmark for assessing whether the data we recorded from each brain area were representative, we compared the amount of visual modulation present in each brain area, at each transformation, with the following rationale. First, all the visual information contained in IT is thought to arrive there after first travelling through V4 (Felleman and Van Essen 1991), and consequently, samples of V4 and IT are comparable only if visual information is equal or higher in the V4 sample. Second, by comparing visual information at each transformation separately, we circumvent issues related to the differences in the format of visual information between the two brain areas described in Fig 2a.

To compare the amounts of visual information in our recorded V4 and IT populations, we computed a single-unit measure of visual modulation described by (Pagan and Rust 2014b) and included in a number of our earlier reports (Pagan et al. 2013; Pagan and Rust 2014a; Roth and Rust 2018a). The advantage of this measure is that it disentangles modulations due to changes in visual identity from other factors, such as top-down target modulation, whereas many other traditional measures (e.g. single-neuron d’ or ROC) do not. More specifically, while measures like single-unit d’ are in fact proportional to modulation by the variable of interest (e.g. visual modulation), they are also inversely proportional to other types of modulation (e.g. target modulation), when they exist, because these other types of modulation act as a form of noise or “nuisance variability” that lowers d’ (Pagan and Rust 2014b). Consequently, for a given value of single-neuron d’, it remains ambiguous whether that value arose from a particular amount of visual modulation in the absence of other modulation types or by larger visual modulation in the presence of other modulation types, and this disambiguation has important consequences for comparing the magnitudes of visual modulation in our recorded V4 and IT data. To resolve this ambiguity, our measure applies an extension of an ANOVA, and like the ANOVA, parses a unit’s total response variance into that which can be attributed to modulation by each type of stimulus variable (e.g. changes in the visual image, changes in target identity, etc.), as well as their nonlinear interactions. We also apply a bias-correction procedure to correct for the over-estimation of these variances due to limited samples, developed and tested extensively in simulation (Pagan and Rust 2014b). Finally, because variance non-intuitively scales as the square of response changes (i.e. when firing rates double, variances quadruple), we calculate modulation in units of standard deviation, computed as the square root of the bias-corrected variance quantities. In sum, this measure of visual modulation quantifies the modulation in a unit’s spike count that can be attributed to changes in the identity of the object in view, and it is computed separately for each of the 5 transformations.

For three of the five transformations (‘left’, ‘small’, ‘up’), mean visual modulation was statistically indistinguishable between V4 and IT (Fig 6a-c). For one transformation (‘big’; Fig 6d) mean visual modulation was larger in V4, but we retained this transformation for subsequent analyses because its incorporation reflected a worst-case scenario against the sampling problem of concern (i.e. one in which V4 has been inadequately sampled). In contrast, for the final transformation (‘right’; Fig 6e), the V4 population had significantly lower performance than IT (p < 1e10^−5^), and investigation of the recorded receptive field locations (Fig 4b) revealed that this was likely due to incomplete sampling at that location. As such, we disregarded this transformation from further analyses. Subsequent analyses are focused on the 4 of 5 transformations in which visual modulation, averaged across transformations, was not statistically distinguishable in V4 as compared to IT, either in the pooled data or in either monkey (Fig 6f; Monkey 1: V4 mean = 0.30, IT mean = 0.27, p = 0.07; Monkey 2: V4 mean = 0.18, IT mean = 0.19, p = 0.93). The fact that visual modulation is matched between V4 and IT across these four transformations suggests that the two populations can and should be compared with approximately matched numbers of units, consistent with previous reports (Rust and DiCarlo 2010).

#### Object identity is more linearly formatted in IT than V4 during the IDMS task

As depicted in Figure 2, information about object identity, across changes in identity-preserving transformations, is reported to be more accessible to a linear read-out in IT as compared to V4 (Rust and DiCarlo 2010). To determine whether this difference between V4 and IT was reflected during the IDMS task, we measured the ability of a 4-way linear object identity classifier, trained at each transformation, to generalize to other transformations. Specifically, “reference performance” was measured as cross-validated classifier performance when the training and testing trials came from the same transformation. “Generalization performance” was measured as cross-validated classifier performance when the testing trials came from the three transformations that were not used for training. To avoid confounding visual and target match modulation, each type of performance was computed separately for target matches and distractors (in all possible combinations) and then averaged (see Methods). Finally, “generalization capacity” was measured as the ratio of generalization over reference performance after subtracting the value expected by chance (where chance = 25%).

Fig 7a depicts how reference and generalization performance grew as a function of population size in each brain area. In V4, generalization performance remained modest across all population sizes whereas V4 reference performance grew at a faster rate. In IT, both reference and generalization performance grew at non-negligible rates. Fig 7b summarizes the results in the two brain areas by plotting the endpoints of the plots in Fig 7a. Generalization capacity, computed as the ratio of generalization over reference performance, was higher in IT as compared to V4 (V4 = 0.16; IT = 0.46; p < 0.001), consistent with IT reflecting a more linearly-separable object representation. This plot also reveals slightly lower reference performance in V4 for matched numbers of units (Fig 7b) despite the two populations reflecting matched average single-unit visual modulation (Fig 6f). We have determined that this small difference can be attributed to the slightly higher variance-to-mean ratio in V4 as compared to IT (reported above, mean Fano factor V4 = 1.56; mean Fano factor IT = 1.46), as opposed to other factors such how the information is tiled across the stimulus space or differences in task-relevant modulation (not shown).

To confirm that IT generalization capacity remained higher even under conditions in which more total visual information was available in V4, we also computed generalization capacity for the full V4 population (n = 650 units). As shown in Figure 7c, generalization capacity remained higher in IT even under these conditions (mean V4 = 0.19; mean IT = 0.46; p < 0.001). Higher generalization capacity also held for each of the transformations individually (Fig 7d; ‘Big’ p < 0.001; ‘Left’ p < 0.001; ‘Small’ p = 0.048; ‘Up’ p < 0.001).

#### Visual representations in V 4 and IT during the IDMS task, summarized

The results presented thus far demonstrate that, consistent with earlier reports, V4 and IT can be compared with approximately matched numbers of units, and that visual representations of object identity are more accessible to a linear population read-out in IT during the IDMS task. As explained above, the rationale behind this experiment included seemingly optimizing the task for top-down integration preferentially in IT over V4 by exploiting differences in how V4 and IT represent object identity invariant to identity-preserving transformations (Fig 2). The results presented thus far suggest that this goal was realized insofar as the visual representations of object identity were in fact measurably different in V4 and IT during this task. In question is whether these differences corresponded with differences in the amount and/or format of non-visual, task-relevant information reflected in the two brain areas.

### Population-based comparisons of top-down modulation in V4 and IT during the IDMS task

The IDMS task involved the monkeys viewing images and determining whether they were target matches (requiring a saccade) or distractors (requiring fixation), and thus can be re-conceptualized as a two-way classification of the same images presented as target matches versus distractors. When applied to neural data, the information available for this task could be could be linearly formatted (Fig 8a) or it could be nonlinear (Fig 8c). Under the assumption that visual modulations are larger than top-down modulations (verified below), top-down modulations serve as the rate-limiting factor for target match task performance, and task performance can, in turn, used as a proxy for comparing the magnitudes of top-down modulation in V4 and IT. To compare the amounts of linearly and nonlinearly formatted information available in V4 and IT during the IDMS task, we compared the performance of a handful of weighted linear and nonlinear decoders applied to each population. By weighting each neuron (e.g. proportional to the amount of task-relevant information that it carries), this process ensures that a unit’s responses were not considered in situations where that unit was not informative. Additionally, we applied the decoders to the neural responses to each transformation separately and then averaged decoder performance across transformations to account for differences in the format of visual information between V4 and IT (Figs 2, 7).

#### Linear target match information, uniform sampling

To quantify the amount of linearly separable target match information in V4 and IT, we computed the cross-validated performance of a Fisher Linear Discriminant, which weights each unit proportional to its d’ for this task, adjusted for any correlations that exist between units (see Methods). To determine whether weighted, uniform sampling of V4 could account for target match information in IT, we randomly selected IT units up to the total numbers of units that we recorded (Fig 8b, gray), and compared this to a random selection of V4 units for matched sized populations (and thus always a subset of the V4 data Fig 8b, red). Cross-validated population performance was higher than chance in V4, but was significantly higher in IT as compared to V4 in both monkeys (Fig 8b, gray versus red). We also compared population performance when all V4 units were included for each monkey (∼3x V4 as compared to IT; Fig 8b, red) and found that performance in V4 remained considerably lower than IT. These results suggest that IT target match information is not directly inherited from V4 under the assumption of a uniform sampling of V4 by IT.

#### Linear target match information, best unit sampling

Evidence from other studies suggests that the brain can learn to preferentially read-out the subset of neurons that carry the most task-relevant information with extensive training (Law and Gold 2009) and the monkeys involved in these experiments were trained extensively. Could a version of the feed-forward proposal in which IT preferentially samples the “best” V4 neurons account for our data? To allow us to address this question, we sampled 3-fold more units in V4 as compared to IT, consistent with anatomical estimates of the ratios of neurons between the two brain areas (DiCarlo et al.2012). To assess whether a “best unit” sampling description of V4 by IT could account for our data, we recomputed performance for V4 and IT populations that were matched in size, but when only the N top-ranked V4 units were included for different sized N. Notably, this analysis differs from the analysis presented above in which all V4 units were included insofar as the computation and assignment of weights based on limited samples is a process that contains some noise, and in scenarios where only a subset of units carry a signal, performance is expected to increase until all signal-carrying units are included, but then can fall slightly with the inclusion of units that do not reflect any signal. In this analysis, units were ranked based on the training data before computing cross-validated performance. We found that V4 performance was slightly higher for the best units as compared to randomly selected units (Fig 8b, cyan vs. red), however, performance for the best V4 units remained lower than IT performance in both monkeys (Fig 8b, cyan vs. gray, p<0.001). These results suggest that during IDMS, IT target match modulation cannot be accounted for via feed-forward propagation of this modulation from V4, even if IT were to sample from the “best” V4 subset.

#### Nonlinear target match information

The results above suggest that IT does not inherit its top-down information from V4 via linear computation (either by sampling uniformly or preferentially sampling from the best V4 units). However, it could be the case that some of the information differentiating target matches and distractors was present at the level of V4, but in nonlinear format (Fig 8c). To quantify the “total” target match information in each brain area, regardless of its format, we measured cross-validated performance for a maximum likelihood (as opposed to linear) classifier (see Methods). Cross-validated population performance was higher than chance in V4 (Fig 8d, red; in both monkeys, compared at n = 108 in monkey 1 and n = 96 in monkey 2, p<0.001), but was also higher in IT as compared to V4 (Fig 8d, gray; in both monkeys, compared at n = 108 in monkey 1, p <0.001 and n = 96 in monkey 2, p=0.009).

#### Population-based comparisons, summarized

Taken together, these results rule out all variants of the “IT: Inherited” proposal presented in Fig 1a, including descriptions in which IT preferentially samples from the best V4 units (Fig 8b, cyan vs gray), as well as descriptions that allow for nonlinear computation on the information arriving in IT from V4 (Fig 8d). Because these results thus suggest that IT target match information is not exclusively inherited via feed-forward projections arriving from V4, we can conclude that, as suggested by the “IT: Integrated” proposal, at least some component of top-down information is integrated directly in IT during the IDMS task (Fig 1b).

### Single-unit comparisons of top-down modulation in V4 and IT during the IDMS task

As a complementary set of analyses, we focused on quantifying how top-down modulation was reflected in the responses of individual units, both as a function to time relative to the onset of the stimulus-evoked response, and as a function of visual tuning.

#### Comparing different types of task-relevant signals in V4 and IT

To interpret the different types of signals that might be reflected in V4 and IT during object search, it is useful to conceptualize how signals that differentiate between target matches versus distractors (“target match signals”) – which reflect the solution to the task – might be computed. When considered in terms of a single 4×4 “looking at” vs. “looking for” matrix (Fig 9a, left), target match signals are reflected as diagonal structure (i.e. you are looking at the same image you are looking for, Fig 9a, right, ‘Target match (four objects)’). In the most straightforward description of target match computation, congruent ‘visual’ information (what you are “looking at”; vertical structure) and ‘target identity’ information (what you are “looking for”; horizontal structure) combine in a nonlinear fashion to compute target match detectors that are selective for one object presented as a target match (‘Target match (one object)’). Finally, these are pooled across the four different objects to create ‘Four object target match detectors’ that respond whenever a target is in view (Fig 9a, right). We found examples of these types of idealized units in V4 and/or IT (Figures 9b-c). In both areas, we found ‘purely visual’ units that responded selectively to images but were not modulated by other factors, such as target identity or whether an image was presented as a target match (Fig 9b-c, ‘Visual’). In contrast, one notable difference between V4 and IT was the existence of a handful of IT units (∼10/204) that reflected the remarkable property of responding to nearly every image presented as a target match (every object at every transformation) but not when those same images were presented as distractors (Fig 9c, ‘Target match (four object)’). We did not find any such units in V4. However, in both V4 and IT, we found units that responded preferentially to individual objects presented as target matches as compared to distractors (Fig 9b-c, ‘Target match (one object)’). We note that while these illustrative examples were chosen because they reflect intuitive forms of pure selectivity, many (if not most) units tended to reflect less intuitive mixtures of visual and task-relevant modulation.

**Figure 8.**
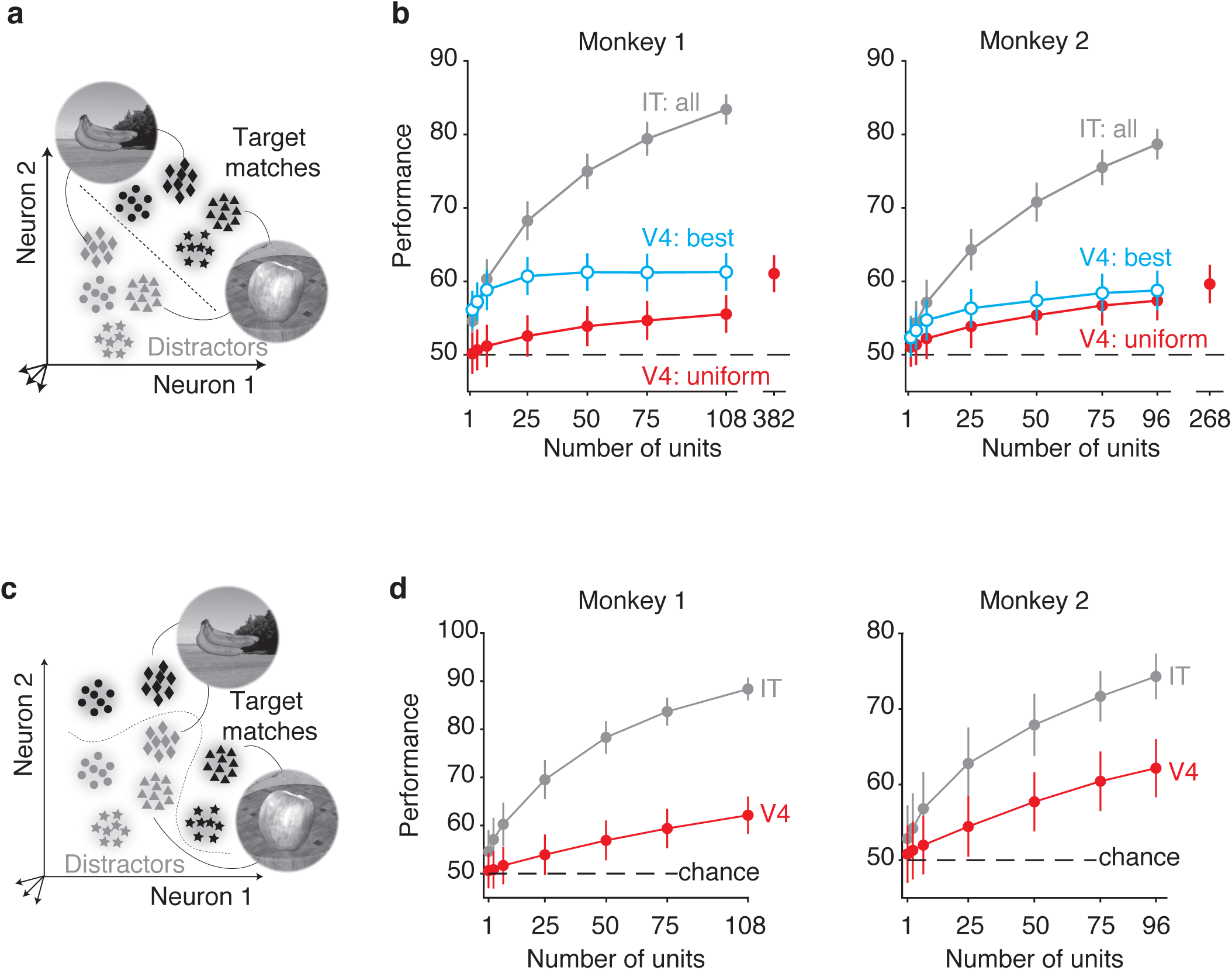
Population-based comparisons of top-down information in V4 and IT. **a)** The IDMS task is presented as a two-way classification of the same images presented as target matches versus as distractors with a linear decision boundary. Each point depicts a hypothetical population response for a population of two neurons on a single trial, and clusters of points depict the dispersion of responses across repeated trials for the same condition. Included are the hypothetical responses to the same images presented as target matches (black) and as distractors (gray). **b)** Performance of a linear classifier trained to classify whether an object was a target match or a distractor, invariant of object identity (at each transformation and averaged across transformations). Red: random subsampling of V4 units; cyan: sampling the best N V4 units based on the training data; gray: random sampling of IT units. Total numbers of IT units: monkey 1: n = 108 units, monkey 2: n = 96 units. Total numbers of V4 units: monkey 1: n = 382, monkey 2: n = 268. Error bars (standard error) reflect the variability that can be attributed to the specific subset of trials chosen for training and testing, and, for subsets of units smaller than the full population, the specific subset of units chosen. Dashed line indicates chance performance. **c)** The IDMS task as in panel a, but formatted nonlinearly. **d)** Performance of a nonlinear, maximum likelihood classifier trained to classify whether an object was a target match or a distractor, invariant of object identity. Performance was assessed at each identity-preserving transformation and then averaged. Error bars (standard error) reflect the variability that can be attributed to the specific subset of trials chosen for training and testing, and, for subsets of units smaller than the full population, the specific subset of units chosen. Dashed line indicates chance performance.

**Figure 9.**
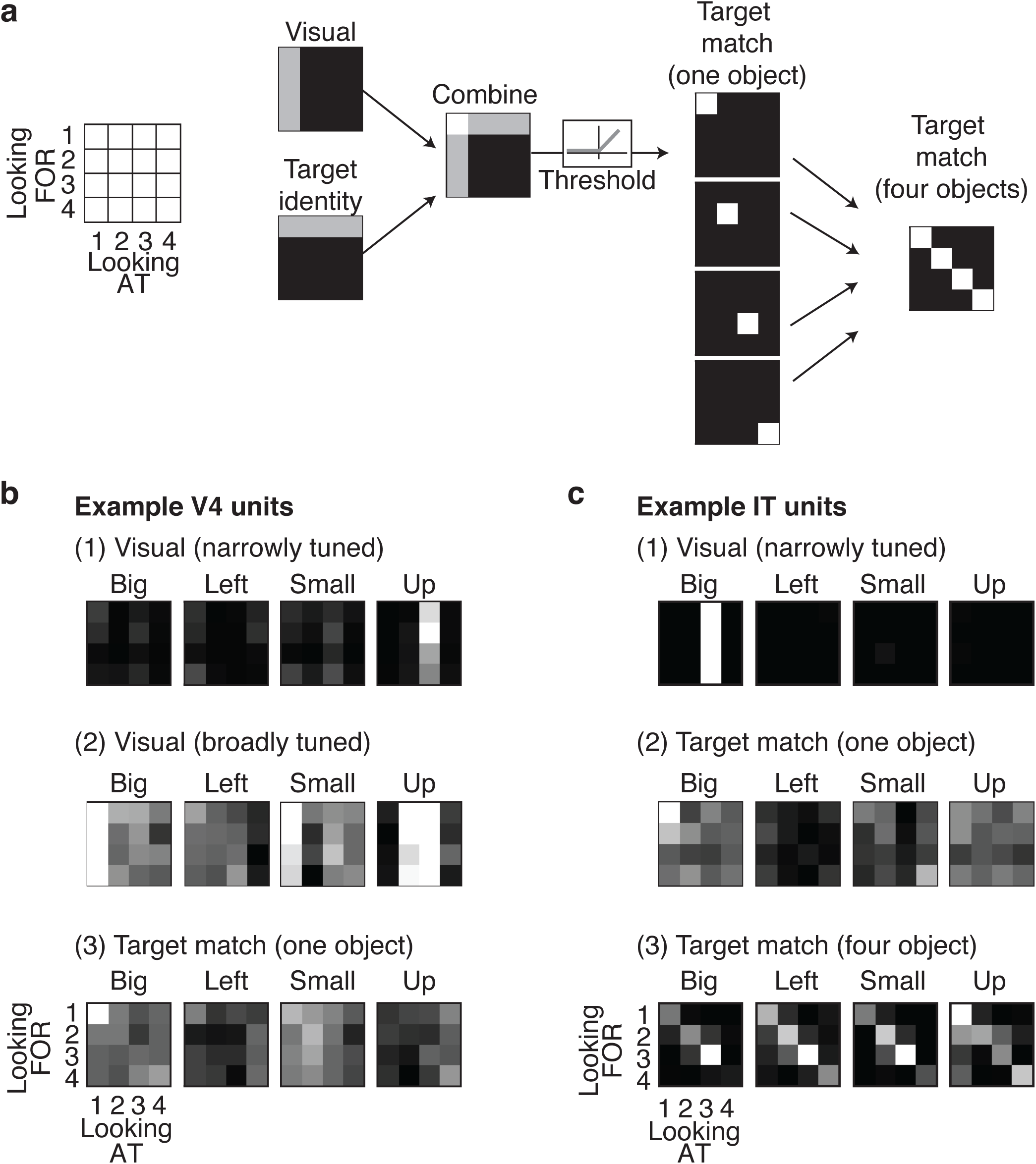
Conceptualizing IDMS task computation. **a)** An idealized depiction of how target match signals, which reflect the solution to the IDMS task, might be computed. For simplicity, the computation is described for one 4×4 slice of the experimental design matrix, which corresponds to viewing each of four objects (‘Looking AT’) in the context of each of four objects as a target (‘Looking FOR’) at one transformation. In the first stage of this idealization of target match computation, a unit reflecting visual information and a unit reflecting persistent target identity information (i.e. working memory) are combined, and the result is passed through a threshold. The resulting unit reflects target match information for one object. Next, four of these units (each with a different object preference) are linearly combined to produce a unit that signals whether a target match is present, regardless of the identity of the object. **b-c)** Example single-unit responses. Responses matrices are plotted as four 4×4 slices of the experimental design matrix. Each slice corresponds to viewing each of four objects (‘Looking AT’) in the context of each of four objects as a target (‘Looking FOR’) at one of the four transformations used. Shown are the response matrices corresponding to 3 example units from V4 and IT. Response matrices were plotted as the average firing rates across trials, and rescaled from the minimum (black) to maximum (white) response across all experimental conditions.

**Figure 10.**
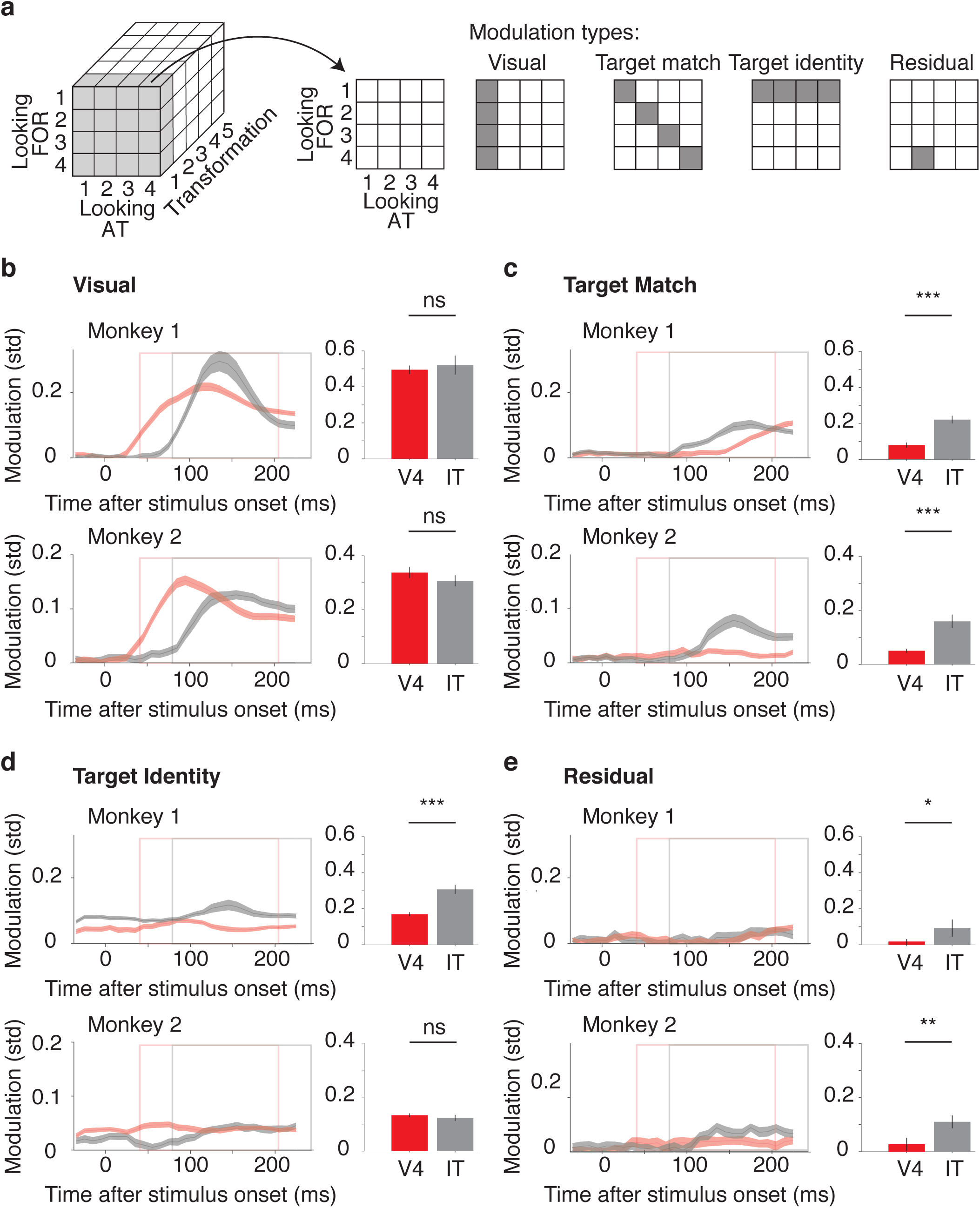
Evolution of different types of single unit modulations in V4 and IT. **a)** To illustrate the different types of task-relevant signals that could be present in V4 and IT, shown is a slice through the IDMS experimental design (Figure 3b), corresponding to one transformation. Shown are visual modulations, which differentiate between different objects in view (vertical structure); target identity modulations, which differentiate between different target objects (horizontal structure); target match modulations, which differentiate between whether objects appear as a target match versus a distractor (diagonal structure); and residual modulations, which differentiate between any other types of conditions (e.g. a response to a particular distractor condition such as looking for object 4 when looking at object 2). **b-e)** Modulations were computed for each type of experimental parameter in units of the standard deviations around each unit’s grand mean spike count (see Results). In each panel, average modulation magnitudes across units in V4 (red; n = 650) and IT (gray; n = 204) shown on the left as a function of time (ms after stimulus onset). Modulation magnitudes, computed in spike count bins 50 ms wide and shifted by 10 ms, are plotted corresponding to the midpoint of each bin, and consequently, end 25 ms before the termination of the last bin. The bar plots show average signal magnitudes quantified within broader spike counting windows indicated by the rectangles on the left (V4: 40-210 ms, red rectangle; IT: 80-250 ms, gray rectangle). Modulation in broader bins can loosely be envisioned as the integral of modulation in narrower bins, however, due to bias correction, this is not exactly the case (see Methods). Single asterisks denote p < 0.05; double asterisks denote p < 0.01; triple asterisks denote p < 0.001; ‘ns’ indicates p > 0.05. Error bars reflect the standard error of modulation across units, computed via a bootstrap procedure.

**Figure 11.**
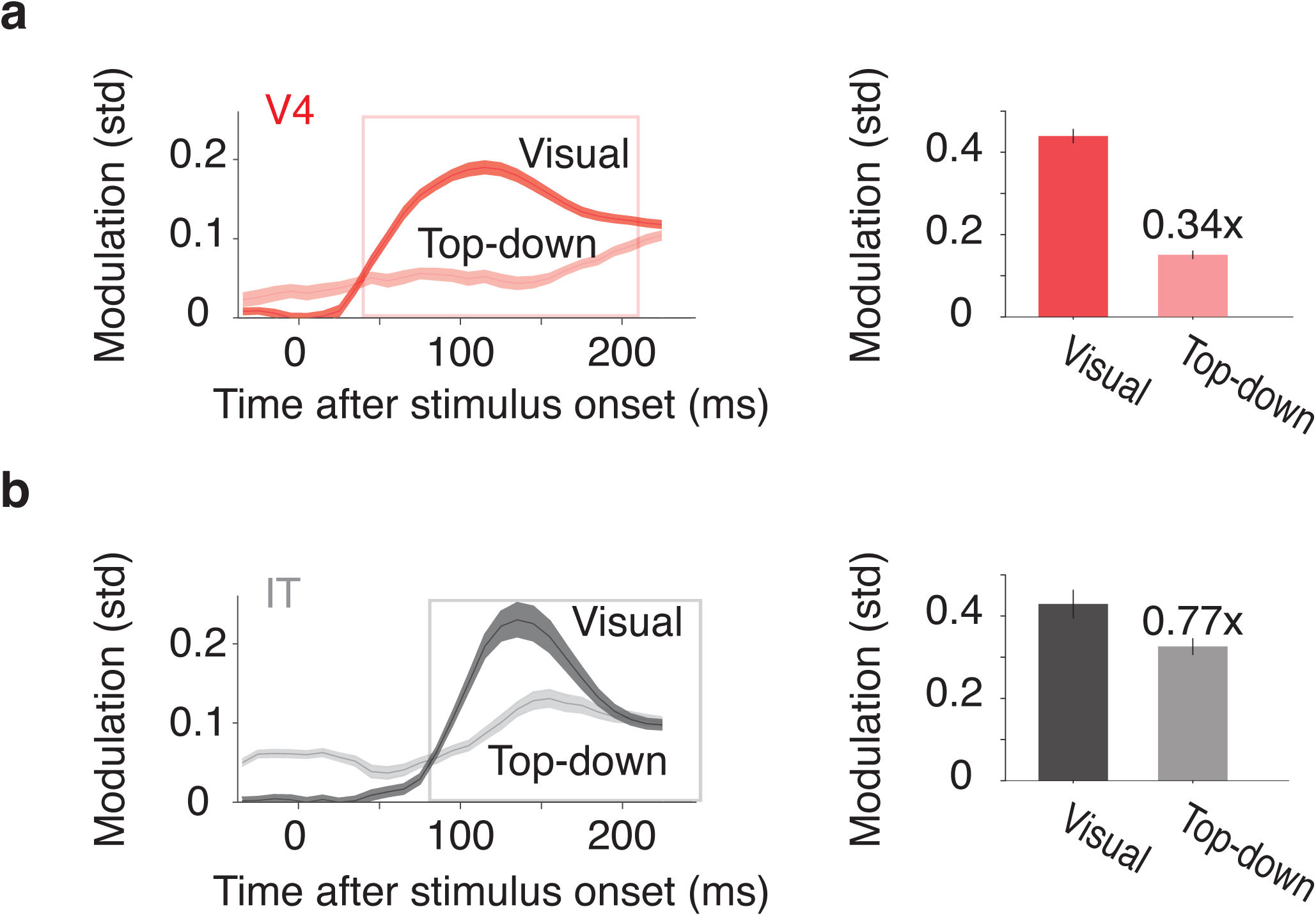
Top down modulations in V4 and IT cortex. **a-b)** Top-down modulations (V4: light red; IT: light gray) were computed as the sum of target identity, target match, and residual modulations, and are shown alongside visual modulations (V4: dark red; IT: dark gray). Mean modulation magnitudes are computed in the same manner and shown with the same conventions as Fig 10. Labels in the bar plots above the top-down modulation magnitudes indicate the proportional size of cognitive relative to visual modulations in each brain area.

To quantify the magnitudes of these different types of task relevant signals in V4 and IT, we extended the procedure presented in Fig 6 to not only quantify ‘visual’ modulation (i.e. modulation that can be attributed to changes in the identity of the visual image), but also other types of non-overlapping modulations that could be attributed to: ‘target identity’ modulation - changes in the identity of a sought target; ‘target match’ modulation - changes in whether an image was a target match or a distractor; and ‘residual’ modulation - nonlinear interactions between visual and target identity that are not target match modulation (e.g. an enhanced response to a particular distractor condition). When considered in terms of a single 4×4 “looking at” vs. “looking for” slice of the experimental design matrix (Fig 3c), these modulations produce vertical, horizontal, diagonal, and off-diagonal structure, respectively (Fig 10a). Notably, this analysis defines target match modulation as a differential response to the same images presented as target matches versus distractors, or equivalently, diagonal structure in the transformation slices presented in Fig 10a. Consequently, units similar to both the ‘target match (one object)’ unit as well as the ‘target match (four object)’ unit (Fig 9a) reflect target match modulation, as both units have a diagonal component to their responses. What differentiates these two types of units is that the ‘Target match (one object)’ unit also reflects selectivity for image and target identity, which is reflected in this analysis as a mixture of target match, visual, and target identity modulation.

We applied this analysis to spike count windows positioned at sliding locations relative to stimulus onset, as well as the same counting windows described for Fig 5 (170 ms; V4: 40-210 ms; IT: 80-250 ms; Fig 7b-e). As expected, visual modulation did not exist before stimulus onset, and visual signals arrived in V4 ∼ 40 ms earlier than in IT in both animals (Fig 10b). In contrast, modulations reflecting information about whether an image was a target match or a distractor (‘target match’ modulation) were considerably smaller in V4 as compared to IT in both animals (Fig 10c; monkey 1 p < 0.001; monkey 2 p < 0.001). In monkey 1, V4 target match modulations increased throughout the viewing period, and reached levels that were similar to those found in IT, but this rise occurred with a delay in V4 relative to IT. This was not replicated in monkey 2, where target match modulations in V4 were small throughout the viewing period.

Modulations reflecting information about the identity of the target (‘target identity’ modulation) were present in both V4 and IT before stimulus onset (Fig 10d), consistent with persistent working memory signals in both brain areas. These persistent signals were stronger in IT as compared to V4 in monkey 1 (p < 0.001) but comparable in size between V4 and IT in monkey 2 (p = 0.23). Lastly, we found that in both V4 and IT, residual modulation was small relative to the other types of modulations (Fig 10e). Residual modulation was larger in IT than V4 in both monkeys (monkey 1: p = 0.048, monkey 2: p = 0.006). To summarize these results, we found that in both monkeys, visual modulation was matched between V4 and IT whereas target match signals were weaker in V4. We also found persistent target identity signals that were reflected in both areas before and throughout the stimulus-evoked period.

As a complementary analysis, we also quantified the total amount of top-down modulation (combined target match, target identity, and residual modulation), and compared it to the evolution of the visual modulation. In both brain areas, top-down modulation was considerable throughout the analysis window (Fig 11, left). During the latency-corrected stimulus-evoked period, top-down modulations were 34% and 77% the size of the visual modulations in V4 and IT, respectively (Fig 11, right). These results demonstrate that considerable non-visual, task-relevant modulations exist in both brain areas, and they also suggest that these are smaller in V4 as compared to IT.

#### The relationship between the visually-evoked response and target match modulation in individual units

The results presented in Fig 10c establish that during the IDMS task, target match modulation is stronger in IT. How does the relationship between the visually-evoked response and the magnitude of target match modulation manifest in the responses of individual units in each brain area? Conceptually, the increase in target match modulation in IT as compared to V4 could be reflected in a similar manner in the two brain areas, such as multiplicative rescaling, but with a larger multiplicative factor in IT. In this scenario, both brain areas would exhibit tuning for target match modulation that is matched in bandwidth to the visually-evoked tuning for images. Alternatively, increases in target match modulation in IT over V4 could result from similar target match modulation magnitudes for each image at which it appears, but with broader tuning across images in IT. We note that these two possibilities are not mutually exclusive.

To explore these issues, we computed the magnitude of target match modulation as a function of the effectiveness of each image at driving individual units to fire. Specifically, we ranked the responses of each unit to the 16 images (4 objects * 4 transformations; averaged across target matches and distractors), and then examined the responses to target matches and distractors separately, after averaging across all units (Fig 12a). Shown are results computed after subtracting the baseline firing rate from each unit’s responses, where negative values correspond to responses below baseline. In both V4 and IT, target match modulation for the highest ranked images was reflected as target match enhancement, and both brain areas exhibited weak target match suppression for the least effective images (Fig 12a). The net impact of target match modulation (averaged across all ranks) was target match enhancement in both brain areas, with stronger net modulation in IT (the difference in the average response to target matches and distractors, averaged across all ranks, divided by the average response to target matches: V4 = +0.07 spks/s; IT = +0.46 spks/s; Fig 12a).

**Figure 12.**
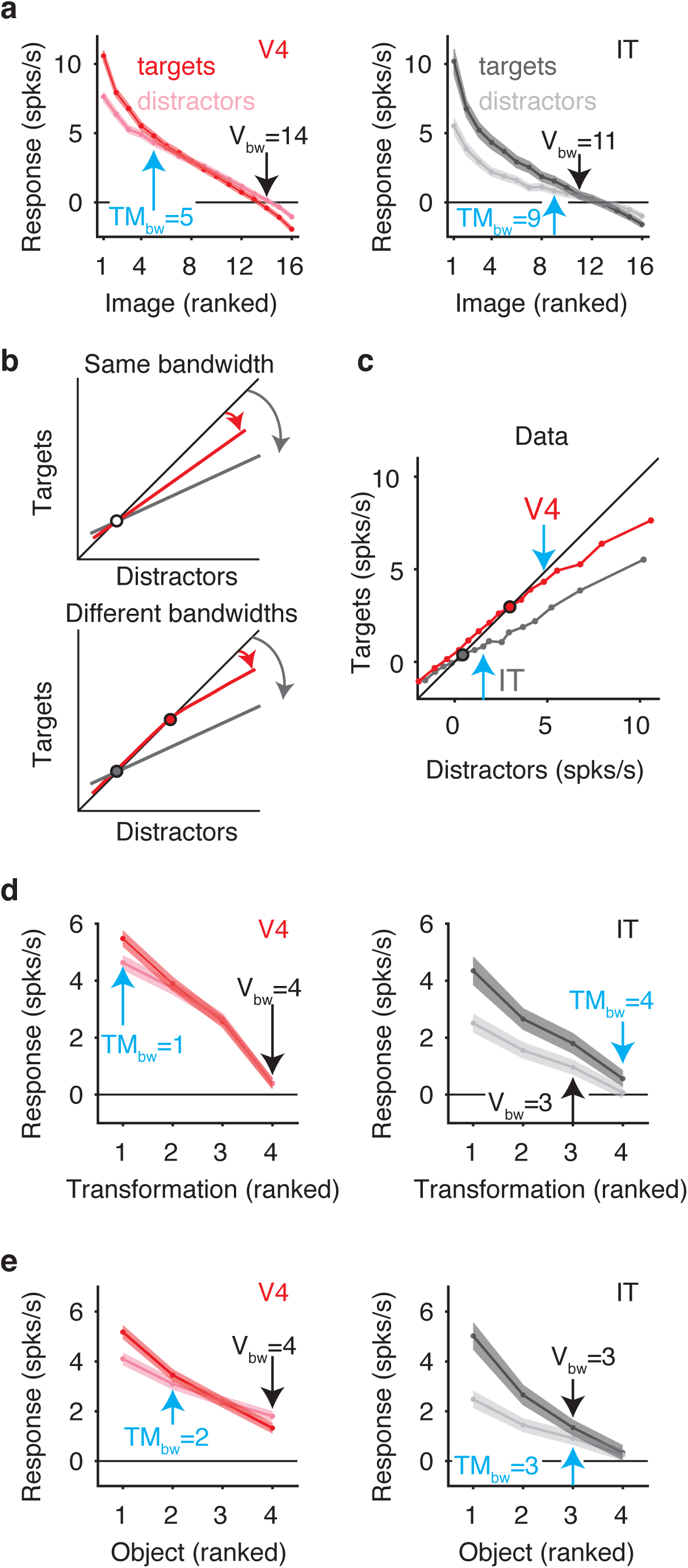
Comparison of the visually-evoked response and target match modulation in individual units. **a)** The spike count responses to target matches (V4: left, red; IT: right, dark gray) and distractors (V4: left, pink; IT: right, light gray) to the 16 different images that made it through the screen presented in Fig 6, ranked by image preference. After subtracting the baseline responses (computed 170 ms before the onset of the first image in each trial), the responses of each unit to each image were ranked by their average response across target matches and distractors. Shown are means and standard errors across all units, after sorting each unit by image rank (V4: n=650; IT: n=204). The bandwidth of the visually-evoked response (V_bw_; black arrows) was computed as the number of images for which the response (averaged across targets and distractors) was significantly larger than baseline (Wilcoxon rank sum test; criterion p = 0.05). Target match bandwidth (TM_bw_; cyan arrows) was computed as the number of images for which the target match response was significantly higher than the distractor response (Wilcoxon rank sum test; criterion p = 0.05). **b)** Schematics showing the hypothetical responses to target matches plotted against hypothetical responses to distractors, similar to a quantile-quantile (Q-Q) plot. Top: shown are two hypothetical populations whose responses are both multiplicatively rescaled but with different factors. This manifests in these plots as changes in the slope of these lines around a fixed pivot point (white dot). Bottom: shown are two hypothetical populations whose responses differ by both their magnitudes of target match modulation as well as the bandwidth of this tuning; changes in bandwidth manifest as different points of intersection with the unity line (red and gray large dots). **c)** V4 and IT spike count responses, plotted as described for panel b. Points of intersection with the unity line are indicated with large red and gray dots and for comparison, the cyan arrows from panel a are included. **d)** Responses to ranked images (averaged across object identity), plotted with the same conventions as panel a. **e)** Responses to ranked objects (averaged across transformation identity), plotted with the same conventions as panel a.

Next we computed tuning bandwidths separately for the visually-evoked response and for target match enhancement. To determine the bandwidth of the visually-evoked response, we determined the number of image ranks with responses significantly above the baseline response (for data averaged across target matches and distractors; Wilcoxon rank sum test; criterion p = 0.05). We found the visually-evoked bandwidths to be slightly broader in V4 (bandwidth= 14 images; Fig 12a left, black arrow) as compared to IT (bandwidth = 11 images; Fig 12a right, black arrow). To determine the bandwidths of target match enhancement, we computed the number of images for which the responses to target matches were significantly higher than the responses to distractors (Wilcoxon rank sum text; criterion p = 0.05). In V4, target match enhancement was more narrowly tuned than the visually-evoked response (target match enhancement bandwidth = 5 images; visually-evoked response bandwidth = 14 images; Fig 12a left, cyan versus black arrows). In IT, these bandwidths were more similar (target match enhancement bandwidth = 9 images; visually-evoked response bandwidth = 11 images; Fig 12a, right; cyan versus black arrows).

To determine whether the observed differences in target match bandwidths between V4 and IT followed simply from target match modulation that was larger in IT, we plotted the responses to target matches against the responses to distractors, similar to a quantile-quantitle or Q-Q plot. In these plots, rescaling the responses by different multiplicative factors but with a fixed bandwidth amounts to changing the slope of the lines around a fixed pivot point (Fig 12b, top) whereas changing the bandwidths results in different points of intersection with the unity line (Fig 12b, bottom). Our data were not well-described as a change in slope around a fixed pivot point, but rather by more narrowly tuned target match enhancement in V4 (Fig 12c). Notably, this narrower bandwidth cannot be explained by a more narrowly tuned visually-evoked response in V4, because the visually-evoked response was slightly broader in V4 as compared to IT (as described above, Fig 12a). These results suggest that increased target match modulation in IT over V4 results from a combination of the two scenarios presented above: it is both larger for the image ranks at which it appears and it is more broadly tuned across images.

The 16 images included in this analysis differed from one another across changes in both object identity as well as object transformation. To determine whether broader target match enhancement in IT as compared to V4 depended differentially on these two factors, we performed the same analysis by ranking each unit per transformation (after averaging across object identity; Fig 12d) or ranking by object identity (after averaging across transformations; Fig 12e). Across transformations, the bandwidth of target match enhancement was considerably narrower in V4 (1 transformation; Fig 12d left, cyan arrow) as compared to IT (4 transformations; Fig 12d right, cyan arrow). Once again, this difference could not be accounted for by a difference in the bandwidth of the visually-evoked response, as this was broader in V4 (4 transformations; Fig 12d left; black arrow) than IT (3 transformations; Fig 12d right; black arrow). Across object identity, the bandwidth of target match enhancement was slightly narrower in V4 (2 objects; Fig 12e left, cyan arrow) as compared to IT (3 objects; Fig 12e right, cyan arrow) whereas the visually-evoked response remained slightly broader in V4 (4 objects; Fig 12e left, black arrow) as compared to IT (3 objects; Fig 12e right, black arrow).

Together, these results suggest that the increase in target match modulation in IT as compared to V4 can be attributed to both an increase in the magnitude of target match modulation for the images that are most effective at driving individual units to fire as well as an increase in bandwidth over the number of images that have this modulation. These IT bandwidth increases, in turn, appear to follow from broader tuning of target match enhancement across both identity-preserving transformations as well as object identity, with a larger influence of the former as compared to the latter.

#### Single-unit comparisons, summarized

Top-down modulations during the IDMS task were larger in IT as compared to V4 (77% vs 34% of the visually-evoked response, respectively), and this could be attributed to larger “target match” and “residual” subtypes of top-down modulation in IT. When examined as a function of the strength of the visually-evoked response, larger IT target match modulation could be attributed to both magnitude as well as bandwidth increases relative to V4.

## DISCUSSION

Finding sought objects requires the brain to compare visual information about the objects in view with information about the currently sought target to compute a signal that reports when a target match has been found. During object search, information about the identity of a sought target and/or whether it is a target match is thought to be fed-back to mid to higher stages of the ventral visual pathway, including V4 and IT, but the specific path this information takes is unclear. In this study, we sought to differentiate between scenarios in which top-down information is integrated directly in IT (Fig 1b) versus those in which it is integrated in V4 and arrives in IT via feed-forward propagation (Fig 1a). We evaluated a number of feed-forward descriptions between V4 and IT, and found none of them could account for the amount of non-visual, task-relevant information present in IT. These included a model in which IT uniformly samples target match signals from V4 (Fig 8b, red), a model in which IT preferentially samples target match signals from the best V4 units (Fig 8b, cyan), and a model that allowed for IT nonlinear processing of inputs arriving from V4 (Fig 8d). Together, these results suggest that during IDMS, top-down, task-specific signals in IT are not exclusively inherited from V4 but rather are integrated within IT, at least in part.

Our study employed a population-based approach in which we recorded neural responses in an unbiased manner while the monkeys performed the task and we counted spikes in the short windows implied by the monkeys’ fast reaction times. Our approaches are a scientific advance over other approaches that artificially increase the signal-to-noise ratio in a dataset in a manner that implicitly makes unrealistic assumptions about the brain. For example, tailoring stimuli to align with the peak of each neuron’s preferences disregards the contributions of neurons that are activated along their curve flank, and counting spikes in long windows that exceed natural reaction times assumes that neural responses are stationary, but they are not (e.g. Pagan and Rust, JNeurosci., 2014). We emphasize that while these types of “increase the SNR approaches” continue to play a fundamental role in the study of visual processing (and we continue to be champions of them in those contexts), the questions under investigation here – which relate to the path of information flow in the brain during the IDMS task – are better studied in a more assumption free manner, using the types of population-based approaches we apply here. In the context of the IDMS task, reaction times are naturally fast, implying spike count windows that are short (e.g. after accounting for latency, 170 ms). In such cases, the brain is expected to operate in a low spike count regime and modulation magnitudes are expected to be small. Similarly, when stimuli are not artificially optimized for each neuron’s preferences, firing rate histograms are expected to have long tails. The way that the brain deals with these types of issues is, of course, by combining the neural responses across many (noisy) neurons via combination rules adjusted for the task at hand (e.g. a weighted linear population read-out). The statistical power in our data arises, similarly, from population-based analyses.

In our study, we were careful to consider whether the larger IT top-down information we observed (relative to V4) was not being confounded with visual information (i.e. stimulus tuning). First, our analyses go to great lengths to demonstrate that larger IT top-down information doe not follow from differences in the total amount of visual information between V4 and IT. Specifically, we demonstrate that all the visual information available in the IT data is also present in randomly sampled populations of a brain area known to provide the input to IT, V4, with approximately matched numbers of units (Fig 6). This establishes that we have recorded from comparable V4 and IT populations and that the visual information in IT can be described as being inherited from V4. If IT also inherited its top-down information from V4, we would expect that when visual information is matched between V4 and IT, top-down information would be matched as well. However, it was not: we found that total top-down information was considerably larger in IT than V4 even when visual information was matched, implying that while visual information can be described as feed-forward, top-down task-relevant information cannot, and thus must be integrated directly within IT itself. It might be of further concern that the larger IT top-down information we observed (relative to V4) erroneously follows from differences in the format of visual information between V4 and IT (e.g. the fact that V4 neurons have smaller receptive fields). We emphasize that our experiment exploited rather than controlled for differences in the format of visual information between V4 and IT, by testing where top-down signals are integrated in a task that required spatial invariance. This experimental design was motivated by findings that top-down effects appear globally across the visual field in V4 despite the small sizes of V4 receptive fields and we thus wondered whether spatially global, top-down integration in V4 could account for top-down signals in IT within a feed-forward framework. A complementary experiment – in which one controls for differences between V4 and IT by matching the task to V4 properties – is interesting and important, but we note that even if that experiment were performed, it would not negate the need to do the one that we did.

We found non-visual, task-specific signals to be sizeable in V4 (∼40% of the size of visual modulation), consistent with many other reports (Moran and Desimone 1985; Haenny et al. 1988; Motter 1994b; Motter 1994a; Luck et al. 1997; McAdams and Maunsell 1999; McAdams and Maunsell 2000; Chelazzi et al. 2001; Ogawa and Komatsu 2004; Bichot et al. 2005; Hayden and Gallant 2005; Mirabella et al. 2007; Cohen and Maunsell 2009; Kosai et al. 2014). At the same time, we also found that non-visual, task-specific modulations to be even larger in IT (∼80% the size of visual modulation). In a previous study, during a visual target search task in which monkeys made a saccade to a target match following the presentation of a sample image, non-visual, task-specific signals were reported to be more similar in V4 and IT (63% and 70% of the visually-evoked response in V4 and IT, respectively; Chelazzi et al. 1998; Chelazzi et al.2001). One notable difference between our study and this earlier work is that our study compared V4 and IT during a version of the delayed-match-to-sample task in which sought target objects could appear at different positions, sizes and background contexts. The fact that top-down, task-specific signals were considerably larger in IT versus V4 in our task may follow from the fact that IT contains a more explicit, linear representation of object identity across these transformations than V4 (reviewed by DiCarlo et al. 2012). Consequently, top-down modulation may be targeted directly to IT in situations that require an invariant object representation whereas the brain might target the pathway differently when tasks have different computational requirements. For example, because V4 receptive fields are smaller and retinotopically organized, V4 might serve as the primary locus for the integration of top-down signals for tasks that require spatial specificity, such as covert spatial attention tasks, and in these tasks little top-down integration might occur in IT (Moran and Desimone 1985). Only one earlier study has reported on the responses of IT neurons in the context of a DMS task in which, objects could appear at different identity-preserving transformations (Leuschow et al. 1994), but this study did not measure signals in V4.

Our results, which demonstrate larger non-visual, task-relevant modulations in IT as compared to V4, are consistent with more general interpretations that the magnitudes of top-down modulation exist in a gradient-like fashion hierarchically along the ventral visual pathway (reviewed by Noudoost et al. 2010). As described above, such gradients are consistent both with the integration of top-down modulation at multiple stages of the pathway (Fig 1b, red) as well as integration at a single locus, followed by feedback within the pathway itself (Fig 1b, cyan). One study (Buffalo et al. 2010) provided evidence supporting the latter description in V1, V2 and V4 in the form of noting that not only the magnitude of modulation was greater in higher visual areas, but it also arrived earlier, consistent with a feed-back description. In our data, this issue was ambiguous: we found that in one monkey, the arrival of target match modulation was delayed in V4 as compared to IT (Fig 10c, Monkey 1) whereas in the other monkey, target match modulation was small in V4 throughout the viewing period (Figure 10c, Monkey 2).

In an earlier series of reports, we compared the responses of IT and its projection area, perirhinal cortex, during a more classic version of the delayed-match-to-sample task (that did not incorporate variation in the objects’ transformations; Pagan et al. 2013; Pagan and Rust 2014a; Pagan et al. 2016). We found that the responses of perirhinal cortex were well-described by a model in which top-down, task-relevant signals were integrated within or before IT consistent with a feed-forward process between IT and perirhinal cortex. The results presented here extend this understanding to suggest that the locus of top-down integration during DMS search tasks is unlikely to exclusively be V4, and that some amount of top-down integration is likely to happen directly within IT itself.

Computing a target match signal requires the combination of the visual representation of the currently viewed scene with a remembered representation of the sought target (e.g. Fig 9a). In an analysis of the same IT data presented here, we found that the IT population misclassified trials on which the monkeys made errors, supporting notions that the IT target match signal is in fact related to the neural signals used to make target match behavioral judgments (Roth and Rust 2018a). The additional target match information present in IT that is not also present in V4 could reflect the implementation of this comparison in IT itself, or alternatively, the comparison might be implemented in a higher order brain area and fed-back to IT cortex. The timing of the arrival of this signal in IT (which peaks at ∼150 ms; Fig 10c) relative to the monkeys’ median reaction times (∼335 ms; Fig 3e), does not rule out the former scenario, but with our data we cannot definitively distinguish between these alternatives. Additionally, in this study monkeys were trained extensively on the images used in these experiments and future experiments will be required to address the degree to which these results hold under more everyday conditions in which monkeys are viewing images and objects for the first time.

In this study, the IDMS task was implemented in short blocks with a fixed target such that a cue image did not need to appear at the beginning of each trial. This design was inspired by studies reporting that when a cue image is presented shortly before a target match (e.g. in the same trial), active target match modulation intermingles with passive, repetition suppression (Miller and Desimone 1994), which we have also found to be true (Pagan et al. 2013). For example, whereas approximately balanced numbers of IT units reflect net target match enhancement versus target match suppression in a classic DMS design (Pagan et al. 2013), IT units nearly universally reflect target match enhancement during the IDMS task (Roth and Rust 2018a). This distinction is important for this study, as our goal was to systematically compare the magnitudes of top-down modulation in V4 and IT whereas repetition suppression is likely to be a largely feed-forward process. At the same time, the block design we employed here also imposes some drawbacks, including less frequent changes of the target identity, and thus less frequent changes of the issuance of new top-down information. Future experiments will be required to assess how these results compare in the context of more dynamic and more realistic object search conditions (where presumably one ceases to look for the same target upon finding it).

## ACKNOWLEDGMENTS

We thank Margot P. Wohl and Krystal Henderson for their technical contributions.

## References

Bichot, N. P., A. F. Rossi and R. Desimone (2005) Parallel and serial neural mechanisms for visual search in macaque area V4. Science 308(5721): 529–534.

Buffalo, E. A., P. Fries, R. Landman, H. Liang and R. Desimone (2010) A backward progression of attentional effects in the ventral stream. Proc Natl Acad Sci U S A 107(1): 361–365.

Chelazzi, L., J. Duncan, E. K. Miller and R. Desimone (1998) Responses of neurons in inferior temporal cortex during memory-guided visual search. J. Neurophysiology 80: 2918–2940.

Chelazzi, L., E. K. Miller, J. Duncan and R. Desimone (1993) A neural basis for visual search in inferior temporal cortex. Nature 363: 345–347.

Chelazzi, L., E. K. Miller, J. Duncan and R. Desimone (2001) Responses of neurons in macaque area V4 during memory-guided visual search. Cereb Cortex 11(8): 761–772.

Cohen, M. R. and J. H. Maunsell (2009) Attention improves performance primarily by reducing interneuronal correlations. Nat Neurosci 12(12): 1594–1600.

Cohen, M. R. and J. H. Maunsell (2011) Using neuronal populations to study the mechanisms underlying spatial and feature attention. Neuron 70(6): 1192–1204.

Desimone, R. and S. J. Schein (1987) Visual properties of neurons in area V4 of the macaque: sensitivity to stimulus form. J Neurophysiol 57(3): 835–868.

DiCarlo, J. J., D. Zoccolan and N. C. Rust (2012) How does the brain solve visual object recognition? Neuron 73(3): 415–434.

Eskandar, E. N., B. J. Richmond and L. M. Optican (1992) Role of inferior temporal neurons in visual memory I. Temporal encoding of information about visual images, recalled images, and behavioral context. Journal of Neurophysiology 68: 1277–1295.

Felleman, D. J. and D. C. Van Essen (1991) Distributed hierarchical processing in the primate cerebral cortex. Cereb Cortex 1(1): 1–47.

Gattass, R., A. P. Sousa and C. G. Gross (1988) Visuotopic organization and extent of V3 and V4 of the macaque. J Neurosci 8(6): 1831–1845.

Gibson, J. R. and J. H. R. Maunsell (1997) The sensory modality specificity of neural activity related to memory in visual cortex. Journal of Neurophysiology 78: 1263–1275.

Haenny, P. E., J. H. Maunsell and P. H. Schiller (1988) State dependent activity in monkey visual cortex. II. Retinal and extraretinal factors in V4. Exp Brain Res 69(2): 245–259.

Hayden, B. Y. and J. L. Gallant (2005) Time course of attention reveals different mechanisms for spatial and feature-based attention in area V4. Neuron 47(5): 637–643.

Hung, C. P., G. Kreiman, T. Poggio and J. J. DiCarlo (2005) Fast readout of object identity from macaque inferior temporal cortex. Science 310(5749): 863–866.

Ito, M., H. Tamura, I. Fujita and K. Tanaka (1995) Size and position invariance of neuronal responses in monkey inferotemporal cortex. J Neurophysiol 73(1): 218–226.

Kosai, Y., Y. El-Shamayleh, A. M. Fyall and A. Pasupathy (2014) The role of visual area V4 in the discrimination of partially occluded shapes. J Neurosci 34(25): 8570–8584.

Law, C. T. and J. I. Gold (2009) Reinforcement learning can account for associative and perceptual learning on a visual-decision task. Nat Neurosci 12(5): 655–663.

Leuschow, A., E. K. Miller and R. Desimone (1994) Inferior temporal mechanisms for invariant object recognition. Cerebral Cortex 5: 523–531.

Li, N., D. D. Cox, D. Zoccolan and J. J. DiCarlo (2009) What response properties do individual neurons need to underlie position and clutter “invariant” object recognition? J Neurophysiol 102(1): 360–376.

Luck, S. J., L. Chelazzi, S. A. Hillyard and R. Desimone (1997) Neural mechanisms of spatial selective attention in areas V1, V2, and V4 of macaque visual cortex. J Neurophysiol 77(1): 24– 42.

Maunsell, J. H., G. Sclar, T. A. Nealey and D. D. DePriest (1991) Extraretinal representations in area V4 in the macaque monkey. Vis Neurosci 7(6): 561–573.

Maunsell, J. H. and S. Treue (2006) Feature-based attention in visual cortex. Trends Neurosci 29(6): 317–322.

McAdams, C. J. and J. H. Maunsell (1999) Effects of attention on orientation-tuning functions of single neurons in macaque cortical area V4. J Neurosci 19(1): 431–441.

McAdams, C. J. and J. H. Maunsell (2000) Attention to both space and feature modulates neuronal responses in macaque area V4. J Neurophysiol 83(3): 1751–1755.

Miller, E. K. and R. Desimone (1994) Parallel neuronal mechanisms for short-term memory. Science 263(5146): 520–522.

Mirabella, G., G. Bertini, I. Samengo, B. E. Kilavik, D. Frilli, C. Della Libera and L. Chelazzi (2007) Neurons in area V4 of the macaque translate attended visual features into behaviorally relevant categories. Neuron 54(2): 303–318.

Moran, J. and R. Desimone (1985) Selective attention gates visual processing in the extrastriate cortex. Science 229(4715): 782–784.

Motter, B. C. (1994a) Neural correlates of attentive selection for color or luminance in extrastriate area V4. J Neurosci 14(4): 2178–2189.

Motter, B. C. (1994b) Neural correlates of feature selective memory and pop-out in extrastriate area V4. J Neurosci 14(4): 2190–2199.

Noudoost, B., M. H. Chang, N. A. Steinmetz and T. Moore (2010) Top-down control of visual attention. Curr Opin Neurobiol 20(2): 183–190.

Ogawa, T. and H. Komatsu (2004) Target selection in area V4 during a multidimensional visual search task. J Neurosci 24(28): 6371–6382.

Op De Beeck, H. and R. Vogels (2000) Spatial sensitivity of macaque inferior temporal neurons. J Comp Neurol 426(4): 505–518.

Pagan, M. and N. C. Rust (2014a) Dynamic target match signals in perirhinal cortex can be explained by instantaneous computations that act on dynamic input from inferotemporal cortex. J Neurosci 34(33): 11067–11084.

Pagan, M. and N. C. Rust (2014b) Quantifying the signals contained in heterogeneous neural responses and determining their relationships with task performance. J Neurophysiol 112(6): 1584–1598.

Pagan, M., E. P. Simoncelli and N. C. Rust (2016) Neural Quadratic Discriminant Analysis: Nonlinear Decoding with V1-Like Computation. Neural Comput: 1–29.

Pagan, M., L. S. Urban, M. P. Wohl and N. C. Rust (2013) Signals in inferotemporal and perirhinal cortex suggest an untangling of visual target information. Nat Neurosci 16(8): 1132– 1139.

Roth, N. and N. C. Rust (2018a) Inferotemporal cortex multiplexes behaviorally-relevant target match signals and visual representations in a manner that minimizes their interference. PLoS ONE In press.

Roth, N. and N. C. Rust (2018b) Rethinking assumptions about how trial and nuisance variability impact neural task performance in a fast processing regime. J Neurophysiol.

Rust, N. C. and J. J. DiCarlo (2010) Selectivity and tolerance (“invariance”) both increase as visual information propagates from cortical area V4 to IT. J Neurosci 30(39): 12978–12995.

Rust, N. C. and J. J. DiCarlo (2012) Balanced increases in selectivity and tolerance produce constant sparseness along the ventral visual stream. J Neurosci 32(30): 10170–10182.

